# Cirrhosis-associated immune dysfunction reflects impaired intestinal CD8αβ⁺ IEL compartmentalization

**DOI:** 10.64898/2026.03.26.713594

**Authors:** Cansu Akkaya, Matthias van Sligtenhorst, Sadia Shaukat, Elodie Modave, Alexander Dumarey, Gabriel Henrique Caxali, Astrid Verbiest, Lise de Meyere, Stiene Vrancken, Lize Van Meerbeeck, Lukas Van Melkebeke, Nina Dedoncker, Stephanie Humblet-Baron, Oliver T. Burton, Adrian Liston, Mitsugu Shimobayashi, Tim Vanuytsel, Schalk van der Merwe, Lidia Yshii, Alexandre Denadai-Souza

## Abstract

Bacterial infections drive decompensation and mortality in cirrhosis, yet the mechanisms underlying cirrhosis-associated immune dysfunction remain unclear. Current models invoke adaptive immune exhaustion, whereas intestinal intraepithelial lymphocytes (IELs), which mediate epithelial surveillance and microbial containment, remain unexplored. We combined spectral flow cytometry of paired duodenal biopsies and blood, plasma proteomics, immunofluorescence and single-cell RNA sequencing in healthy volunteers and patients with compensated or decompensated cirrhosis. The healthy duodenal epithelium was dominated by CD8αβ⁺ IELs, which were depleted in cirrhosis, whereas innate cytotoxic populations expanded. CD8αβ⁺ IEL loss correlated with plasma markers of bacterial translocation. Residual IELs showed attenuated antimicrobial programs, confirmed by reduced granulysin protein, and enriched cytotoxic pathways. IEL depletion was accompanied by reduced CCR9 on circulating and mucosal CD8⁺ T cells and elevated plasma CCL25. Exposing blood mononuclear cells to patient-level CCL25 concentrations reduced surface CCR9, indicating ligand-driven desensitization. In contrast, circulating lymphocytes retained functional competence and lacked exhaustion markers. Cirrhosis-associated immune dysfunction thus reflects failed intestinal compartmentalization of CD8αβ⁺ T cells, linked to CCL25–CCR9 axis disruption, rather than systemic exhaustion.

## Introduction

Cirrhosis represents the final common pathway of chronic liver disease and is characterized by progressive fibrosis, architectural distortion, portal hypertension, and chronic inflammation (Schwabe & Brenner, 2025). Clinically, disease progression spans compensated stages with preserved hepatic function and decompensated stages marked by ascites, encephalopathy, variceal bleeding, and increased short-term mortality (Costa *et al*, 2025). Bacterial infections are among the strongest precipitants of acute decompensation and acute-on-chronic liver failure, profoundly influencing disease progression and survival (Villanueva *et al*, 2021). These observations have placed immune dysfunction at the center of cirrhosis pathophysiology.

Cirrhosis-associated immune dysfunction is traditionally conceptualized as the coexistence of systemic inflammation and immune paralysis (Albillos *et al*, 2014). Current models predominantly emphasize innate immune abnormalities, including excessive inflammatory activation together with impaired antimicrobial responses (Bernsmeier *et al*, 2020). In parallel, adaptive immune dysfunction has frequently been interpreted through the framework of T-cell exhaustion (Lebosse *et al*, 2019; Martin-Mateos *et al*, 2019). However, bona fide exhaustion is defined by sustained inhibitory receptor expression coupled to progressive functional impairment (Barber *et al*, 2006; Blank *et al*, 2019; Wherry, 2011), and whether this state truly characterizes adaptive immunity in cirrhosis remains unresolved. Moreover, most studies have focused on circulating immune cells, despite the fact that infections in cirrhosis predominantly arise from failure to contain intestinal microbes at mucosal barrier surfaces.

Indeed, disruption of intestinal barrier homeostasis and microbial translocation are now recognized as central drivers of cirrhosis progression (Smets *et al*, 2025). The proximal small intestine, particularly the duodenum, appears to represent a critical interface in this process (Maccioni *et al*, 2020). Effective immune surveillance at this site is therefore essential to restrict bacterial expansion and prevent systemic dissemination. Nevertheless, the organization of adaptive immune defense within the intestinal epithelial compartment in cirrhosis remains poorly understood.

Intestinal intraepithelial lymphocytes (IELs) constitute a specialized immune compartment strategically positioned within the epithelial layer to provide immediate barrier surveillance (Hoytema van Konijnenburg *et al*, 2017; Lockhart *et al*, 2024). These cells integrate antigen-specific and innate-like sensing programs to eliminate infected or stressed epithelial cells while preserving tissue integrity. Conventional CD8αβ⁺ IELs are particularly important for epithelial immune surveillance and depend on CCR9-mediated trafficking toward epithelial-derived CCL25 gradients for proper intestinal localization (Stenstad *et al*, 2007). Despite their central role in mucosal host defense, IELs have remained largely unexplored in cirrhosis.

Here, we hypothesized that cirrhosis-associated immune dysfunction does not primarily reflect generalized adaptive immune exhaustion, but rather defective spatial organization of immune responses across compartments. Specifically, we postulated that preserved systemic lymphocyte competence may coexist with impaired deployment of immune effectors at the intestinal epithelial interface. To address this, we performed integrated high-dimensional immune profiling of paired duodenal biopsies and peripheral blood across cirrhosis stages using spectral flow cytometry, plasma proteomics, immunofluorescence, and single-cell RNA sequencing. Our findings identify profound remodeling of the intestinal intraepithelial immune compartment and implicate dysregulation of the CCL25–CCR9 axis as a candidate mechanism underlying defective mucosal immune compartmentalization in cirrhosis.

## Results

### Study cohort and integrated immune profiling strategy

To define how cirrhosis progression reshapes systemic and mucosal immunity, we performed integrated immune profiling of paired peripheral blood and duodenal biopsies obtained from healthy volunteers (HV), patients with compensated cirrhosis (CC), and patients with decompensated cirrhosis (DC). The primary paired cohort comprised 18 individuals, including 6 HV, 6 CC, and 6 DC donors, from whom a total of 108 duodenal biopsies and matched peripheral blood samples were collected for spectral flow cytometry and plasma proteomic analyses. To increase statistical power for systemic immune profiling, peripheral blood samples from four additional donors were included, resulting in 8 HV, 7 CC, and 7 DC samples for circulating analyses. Inclusion of additional donors for protein validation and scRNA-seq resulted in a cohort consisting of 13 HV, 8 CC, and 11 DC donors (Table 1, Appendix Fig. S1).

**Table 1.** Population characteristics, disease parameters and co-morbidities.

| **Population characteristics** | **Healthy volunteer** | **Compensated cirrhosis** | **Decompensated cirrhosis** |
| --- | --- | --- | --- |
| **Samples (N)** | 13 | 8 | 11 |
| **Age median (IQR)** | 36 (28.5 - 59.5) | 65 (63.5 - 69.75) | 62 (57 - 68) |
| **Age range** | 22 - 68 | 54 - 72 | 54 - 75 |
| **Gender (F/M)** | 7/6 | 1/7 | 2/9 |
| **BMI median (IQR)** | 22.6 (20.2 - 25.9) | 26.8 (24.1 - 30.1) | 27.5 (24.9 - 33.4) |
| **BMI range** | 18.4 - 26.9 | 20.6 - 36.4 | 20.8 - 40.9 |
| **MELD median (IQR)** | ND | 8 (6.3 - 10) | 11 (8 - 21) |
| **MELD range** | ND | 6 - 10 | 6 - 24 |
| **Child Pugh (IQR)** | ND | 5 (5 - 5) | 8 (6 - 9) |
| **Child Pugh range** | ND | 5 - 6 | 5 - 11 |

| **Disease parameters** | **Healthy volunteer** | **Compensated**<br>**cirrhosis** | **Decompensated**<br>**cirrhosis** |
| --- | --- | --- | --- |
| **Samples (N)** | 13 | 8 | 11 |
| **Portal hypertension** | 0 (0%) | 5 (62.5%) | 9 (81.8%) |
| **Hepatocellular**<br>**carcinoma** | 0 (0%) | 1 (12.5%) | 1 (9.1%) |
| **Ascites** | 0 (0%) | 0 (0%) | 8 (72.7%) |
| **Hepatic**<br>**encephalopathy** | 0 (0%) | 0 (0%) | 3 (27.3%) |
| **Varices** | 0 (0%) | 3 (37.5%) | 4 (36.4%) |
| **Hepatorenal syndrome** | 0 (0%) | 0 (0%) | 1 (9.1%) |

| **Co-morbidities** | **Healthy volunteer** | **Compensated**<br>**cirrhosis** | **Decompensated**<br>**cirrhosis** |
| --- | --- | --- | --- |
| **Diabetes** | 0 (0%) | 3 (37.5%) | 4 (36.4%) |
| **Arterial hypertension** | 0 (0%) | 5 (62.5%) | 6 (54.5%) |
| **Hypercholesterolemia** | 0 (0%) | 3 (37.5%) | 3 (27.3%) |
| **Chronic kidney disease** | 0 (0%) | 0 (0%) | 1 (9.1%) |

As expected, cirrhosis cohorts displayed higher age and body mass index compared with healthy volunteers, together with progressive increases in MELD and Child–Pugh scores from compensated to decompensated disease. Decompensated cirrhosis was associated with portal hypertension-related complications including ascites, encephalopathy, and varices. To resolve immune remodeling across compartments, we combined high-dimensional spectral flow cytometry, plasma proteomics, immunofluorescence imaging, intracellular cytokine profiling, and single-cell RNA sequencing.

### The duodenal intraepithelial compartment is dominated by CD8αβ^+^ T cells in health

Because the intestinal intraepithelial compartment has never been systematically characterized in cirrhosis, we first established a reference immune landscape in healthy individuals with our 44-color immunophenotyping panel. High-dimensional spectral flow cytometry identified eight major lymphocyte populations within the intraepithelial compartment and matched peripheral blood, including B cells (CD19⁺CD20⁺), natural killer (NK) cells (CD56⁺CD3⁻), NKT cells (CD56⁺CD3⁺), γδ cells (TCRγδ⁺CD3⁻), γδ T cells (TCRγδ⁺CD3⁺), CD4⁺ T cells (TCRαβ⁺CD4⁺), CD4⁺CD8αα⁺ double-positive T cells (TCRαβ⁺CD4⁺CD8α⁺CD8β⁻), and CD8αβ⁺ T cells (TCRαβ⁺CD4⁻CD8α⁺CD8β⁺) (Fig. 1A, Fig. EV1A).

**Fig. 1.**
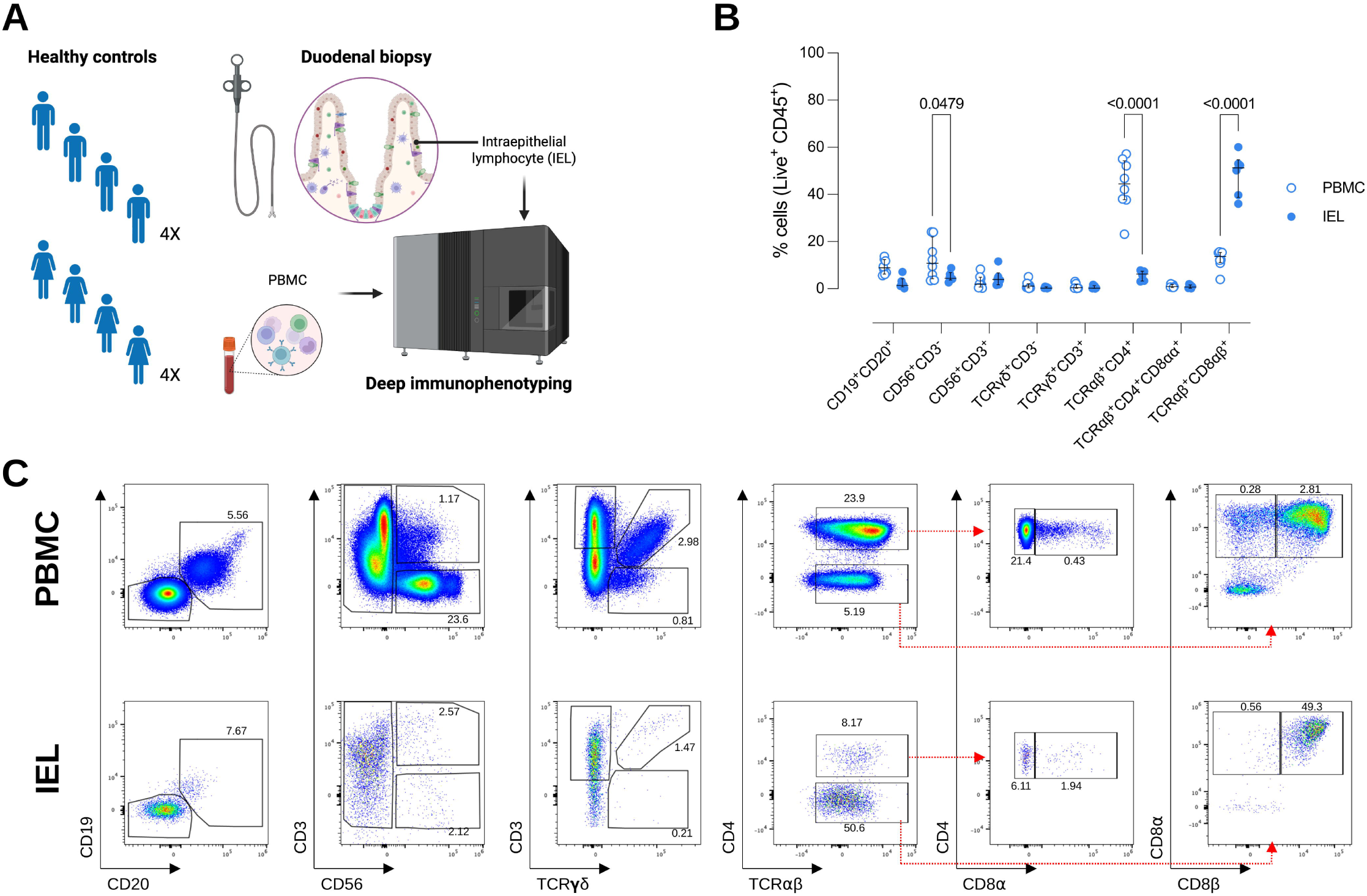
The duodenal intraepithelial compartment is dominated by CD8ab^+^ T cells in health. **(A)** Study design: peripheral blood and duodenal biopsies were obtained from healthy volunteers (HV). Plasma and peripheral blood mononuclear cells (PBMCs) were isolated from whole blood, and intraepithelial lymphocytes (IELs) were dissociated from duodenal biopsies. Immune populations were analyzed by high-dimensional spectral flow cytometry. **(B)** Frequencies of indicated immune cell subsets among live CD45⁺ cells in PBMCs (n = 8) and duodenal IELs (n = 6) from HV. Each dot represents one donor. **(C)** Representative gating of lymphocyte subsets in PBMCs (top) and duodenal IELs (bottom) from HV: CD19⁺CD20⁺ B cells, CD56⁺CD3⁺ NKT cells and CD56⁺CD3⁻ NK cells, TCRγδ⁺ and TCRαβ⁺ T cells, CD4⁺ and CD4⁻ subsets within TCRαβ⁺ T cells, CD8α expression within CD4^+/^⁻ T cells, CD4^+^CD8αα and CD4^-^CD8αβ subsets. Red dashed arrows indicate the sequential gating hierarchy. Numbers indicate the frequency of parent populations. Data are presented as median ± IQR. Statistical analysis was performed with a two-way ANOVA and Bonferroni’s multiple comparison test; p values ≤0.05 are indicated.

Comparative compartmental analysis revealed marked immune specialization between blood and epithelium. Peripheral blood was enriched in NK cells and CD4⁺ T cells, whereas the intraepithelial compartment was dominated by CD8αβ⁺ T cells (Fig. 1B,C), consistent with their established role in epithelial immune surveillance and barrier protection (Jung *et al*, 2025). These data corroborate that the healthy duodenal epithelium is a highly specialized CD8αβ⁺ T-cell niche and provide a baseline for evaluating disease-associated immune remodeling in cirrhosis.

### Cirrhosis reshapes circulating T-cell differentiation without inducing exhaustion

We next investigated whether progressive cirrhosis was associated with major alterations in the circulating lymphocyte compartment (Fig. 2A). The overall abundance of principal circulating lymphocyte populations remained largely preserved across disease stages, with no differences in B-cell, NK-cell, or T-cell subset frequencies between healthy volunteers and cirrhosis cohorts (Fig. 2B).

**Fig. 2.**
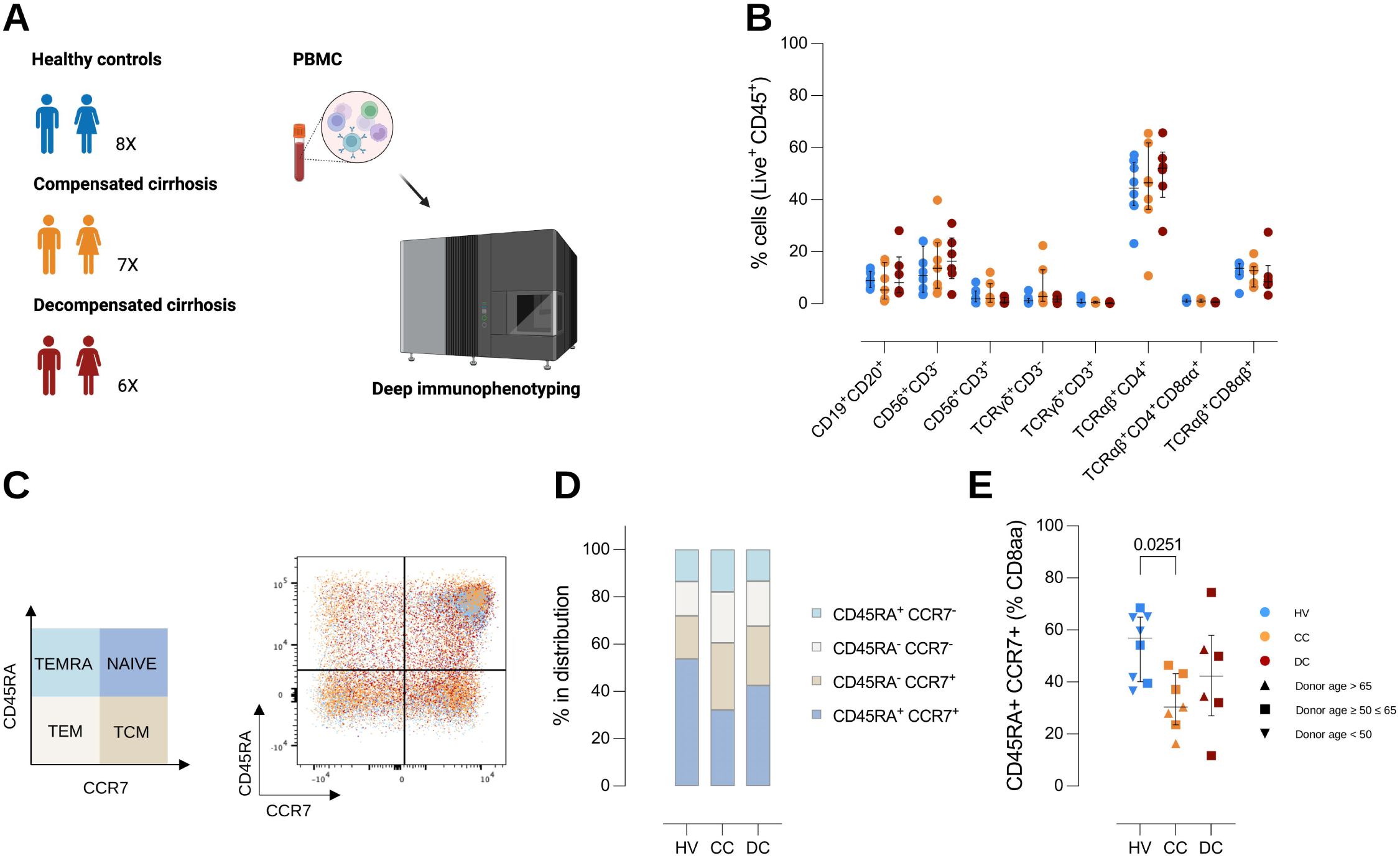
Cirrhosis reshapes the differentiation state of peripheral CD8⁺ T cells. **(A)** Study cohort and workflow: PBMCs from HV, patients with compensated cirrhosis (CC), and decompensated cirrhosis (DC) were subjected to deep immunophenotyping. **(B)** Frequencies of major lymphocyte subsets among live CD45⁺ cells, including CD19⁺CD20⁺ B cells, CD56⁺CD3⁻ NK cells, CD56^+^CD3^+^ NKT cells, and T-cell populations (TCRγδ⁺CD3⁻, TCRγδ⁺CD3⁺, TCRαβ⁺CD4⁺, TCRαβ⁺CD4⁺CD8αα⁺, and TCRαβ⁺CD8αβ⁺) across groups. **(C)** Definition (left) and representative flow cytometry plots (right) of CD8⁺ T cells in PBMC differentiation states based on CD45RA and CCR7 expression: naïve (CD45RA⁺CCR7⁺), central memory (TCM, CD45RA⁻CCR7⁺), effector memory (TEM, CD45RA⁻CCR7⁻), and TEMRA (CD45RA⁺CCR7⁻), overlaid for HV (blue), CC (orange), and DC (red). **(D)** Distribution of TCRαβ⁺CD8αβ⁺ in PBMCs (naïve, TCM, TEM, TEMRA) shown as stacked bar plots for each group. **(E)** Frequency of naïve (CD45RA⁺CCR7⁺) cells within TCRαβ⁺CD8αβ⁺ cells. **(B,E)** Each dot represents one donor from HV (n = 8), CC (n = 7) and DC (n = 6). Data are presented as median ± IQR. Outlier, if present, was removed with Grubbs’ method (alpha = 0.05). Statistical analysis was performed with **(B)** a two-way ANOVA and Bonferroni’s multiple comparison test or **(E)** a Kruskal-Wallis test and Dunn’s multiple comparisons test; comparing CC and DC with HV; p values ≤0.05 are indicated, symbols are coded by donor age.

Despite preserved overall composition, circulating CD8αβ⁺ T cells displayed differentiation remodeling. Stratification according to CCR7 and CD45RA expression demonstrated redistribution from naïve toward effector and memory states in cirrhosis, consistent with chronic inflammatory activation and persistent antigen exposure associated with advanced liver disease (Fig. 2C–E) (Niehaus *et al*, 2025). No significant difference in circulating CD4^+^ T cells or B-cell subsets was observed (Fig. EV1B,C). Thus, cirrhosis primarily reshapes the differentiation landscape of circulating CD8αβ⁺ T cells without inducing major depletion of circulating lymphocyte populations.

Because adaptive immune dysfunction in cirrhosis is frequently interpreted within the framework of T-cell exhaustion, we next assessed expression of canonical inhibitory receptors across compartments. In peripheral blood, there was no increase in PD-1⁺ and CTLA-4⁺ cells across groups in both CD4⁺ and CD8αβ⁺ T cells, rather, there was a significant decrease in PD-1⁺ cells in decompensated cirrhosis (Fig. EV2A,B). Similarly, no significant enrichment of PD-1 or CTLA-4 expression was detected in intraepithelial or lamina propria T-cell compartments (Fig. EV2C–E).

To determine whether preserved phenotypic profiles were accompanied by maintained functional competence, PBMCs were stimulated with CD3/CD28 and subjected to intracellular cytokine profiling and secretome analysis. Stimulation induced robust lineage-specific cytokine responses across all groups (Fig. 3A, Appendix Fig. S2), indicating preserved lymphocyte responsiveness in cirrhosis. CD4⁺FOXP3⁺ cells preferentially induced IL-10, CD4⁺TBET⁺ cells upregulated TNF, CD4⁺RORγT⁺ cells induced IFN-γ, IL-17A, TNF, and granzyme B, whereas CD8⁺ and TCRVα7.2⁺ T cells displayed strong cytotoxic cytokine programs.

**Fig. 3.**
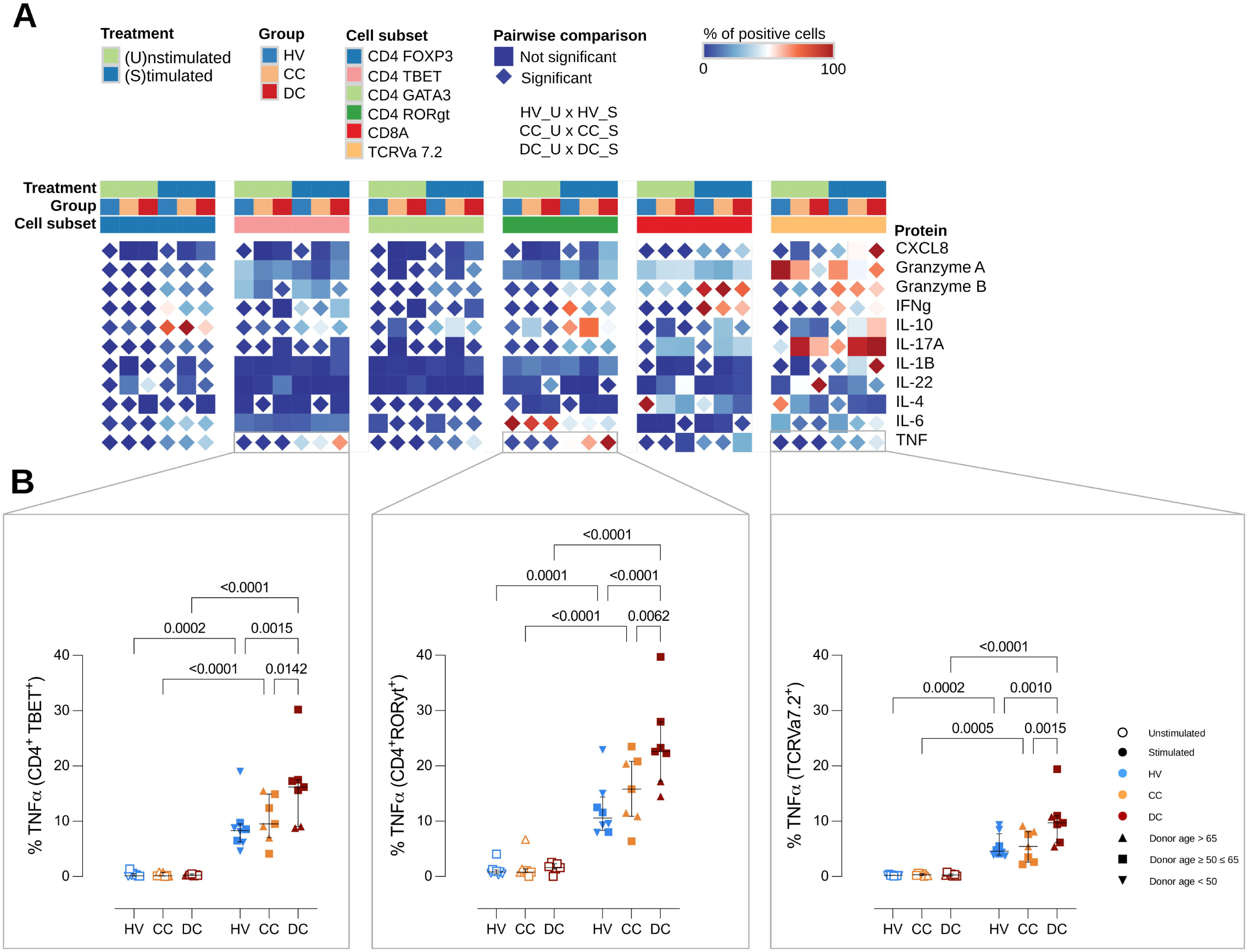
Circulating lymphocytes retain effector competence in cirrhosis. **(A)** Heatmap summarizing intracellular protein production by CD4⁺ T-cell subsets (Th1: CD4⁺TBET⁺, Th2: CD4⁺GATA3⁺, Th17: CD4⁺RORγt⁺, Treg: CD4⁺FOXP3⁺), CD8α⁺ T cells, and TCRVα7.2⁺ T cells from HV, CC, and DC under unstimulated and stimulated conditions. Color intensity indicates the percentage (0 – 100%) of intracellular protein–positive cells; statistically significant pairwise comparisons (p ≤0.05) are denoted by diamond shapes. **(B)** Frequency of TNF-producing T cells following stimulation within Th1 (CD4⁺TBET⁺), Th17 (CD4⁺RORγt⁺), and TCRVα7.2⁺ subsets under unstimulated and stimulated conditions. Each dot represents one donor from HV (n = 8), CC (n = 7) and DC (n = 7). Data are presented as median ± IQR. Statistical analysis was performed with a matched repeated-measures two-way ANOVA and Tukey’s multiple comparisons test; p values ≤0.05 are indicated, symbols are coded by donor age.

Most functional responses were comparable between groups. However, TNF-producing populations were selectively enriched in decompensated cirrhosis, particularly within CD4⁺TBET⁺, CD4⁺RORγT⁺, and TCRVα7.2⁺ T-cell subsets (Fig. 3B). Secreted inflammatory mediators measured in conditioned media further confirmed preservation of T-cell effector capacity across disease stages (Fig. EV3).

Together, these findings indicate that progressive cirrhosis is not characterized by overt adaptive immune paralysis or canonical T-cell exhaustion. Instead, circulating lymphocytes retain functional competence despite progressive inflammatory remodeling.

### Progressive systemic inflammation is associated with early and persistent CCL25 elevation

To define how systemic inflammatory programs evolve across cirrhosis stages, we quantified 89 inflammatory and immune-related plasma proteins using targeted proteomics. Principal component analysis revealed progressive inflammatory remodeling across disease stages, with decompensated cirrhosis clearly separating from healthy volunteers, whereas compensated cirrhosis displayed an intermediate profile (Fig. 4A).

**Fig. 4.**
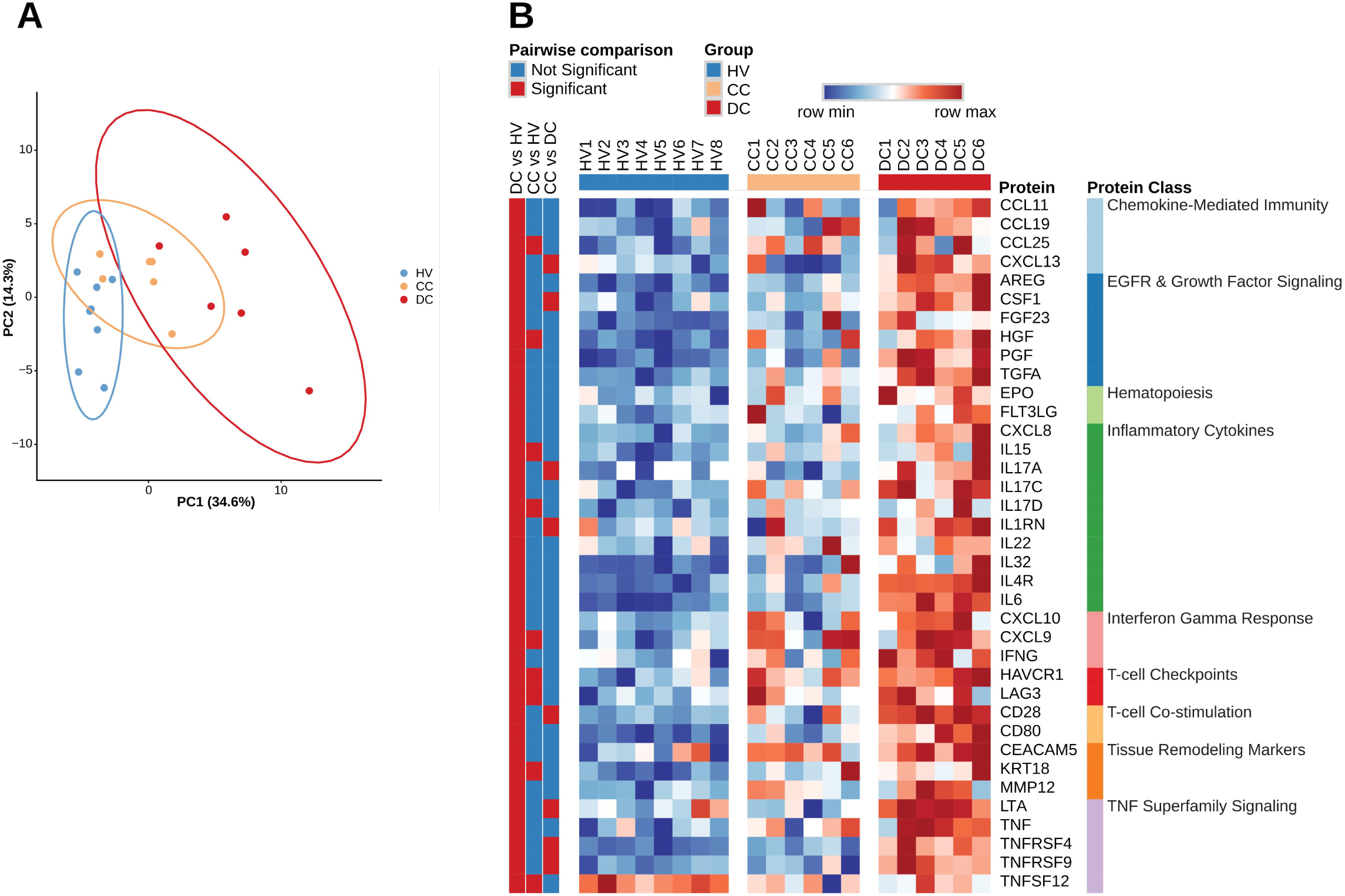
Progressive systemic inflammation is associated with early and persistent CCL25 elevation. **(A)** Principal component analysis (PCA) of inflammation and immune proteins in plasma samples from HV (n = 8), CC (n = 6), and DC (n = 6) patients. **(B)** Heatmap of plasma proteins across HV, CC, and DC, grouped by functional categories. Scaled (z-score) protein levels are displayed, significant pairwise comparisons between groups are indicated. Statistical analysis was performed with a Kruskal-Wallis test and Dunn’s multiple comparisons test; p values ≤0.05 are indicated.

Compensated cirrhosis was associated with relatively limited inflammatory remodeling, with differential expression restricted to nine proteins involved in chemokine-mediated immunity (CCL25), EGFR and growth factor signaling (HGF), inflammation (IL15, IL17D), interferon gamma response (CXCL9), T-cell checkpoints (HAVCR1, LAG3), tissue remodeling (KRT18), and TNF superfamily signaling (TNFSF12).

In contrast, decompensated cirrhosis was associated with an additional differential expression of 28 proteins compared with healthy volunteers, eight of which were also differentially expressed compared with compensated cirrhosis. The increased expression of in total 36 mediators in decompensated cirrhosis were involved in aforementioned processes, as well as hematopoiesis and T-cell co-stimulation, and indicate a broad inflammatory activation. Notably, several TNF-superfamily members were selectively enriched in decompensated cirrhosis compared with both healthy volunteers and compensated disease, consistent with progressive amplification of inflammatory signaling during disease progression (Fig. 4B, Table EV1).

Importantly, CCL25 elevation was observed across both compensated and decompensated disease, whereas the broader inflammatory program emerged predominantly during advanced disease progression. Although the cross-sectional design precludes definitive temporal inference, these observations position CCL25 dysregulation as an early-associated feature of cirrhosis progression rather than a secondary consequence of advanced systemic inflammation.

### The intestinal epithelial immune compartment undergoes marked remodeling in cirrhosis

Having established that systemic lymphocyte competence remains largely preserved, we next investigated whether cirrhosis selectively alters the intestinal intraepithelial immune compartment (Fig 5A). Analysis of intraepithelial lymphocyte composition revealed pronounced remodeling of the duodenal epithelial niche in cirrhosis (Fig. 5B).

**Fig. 5.**
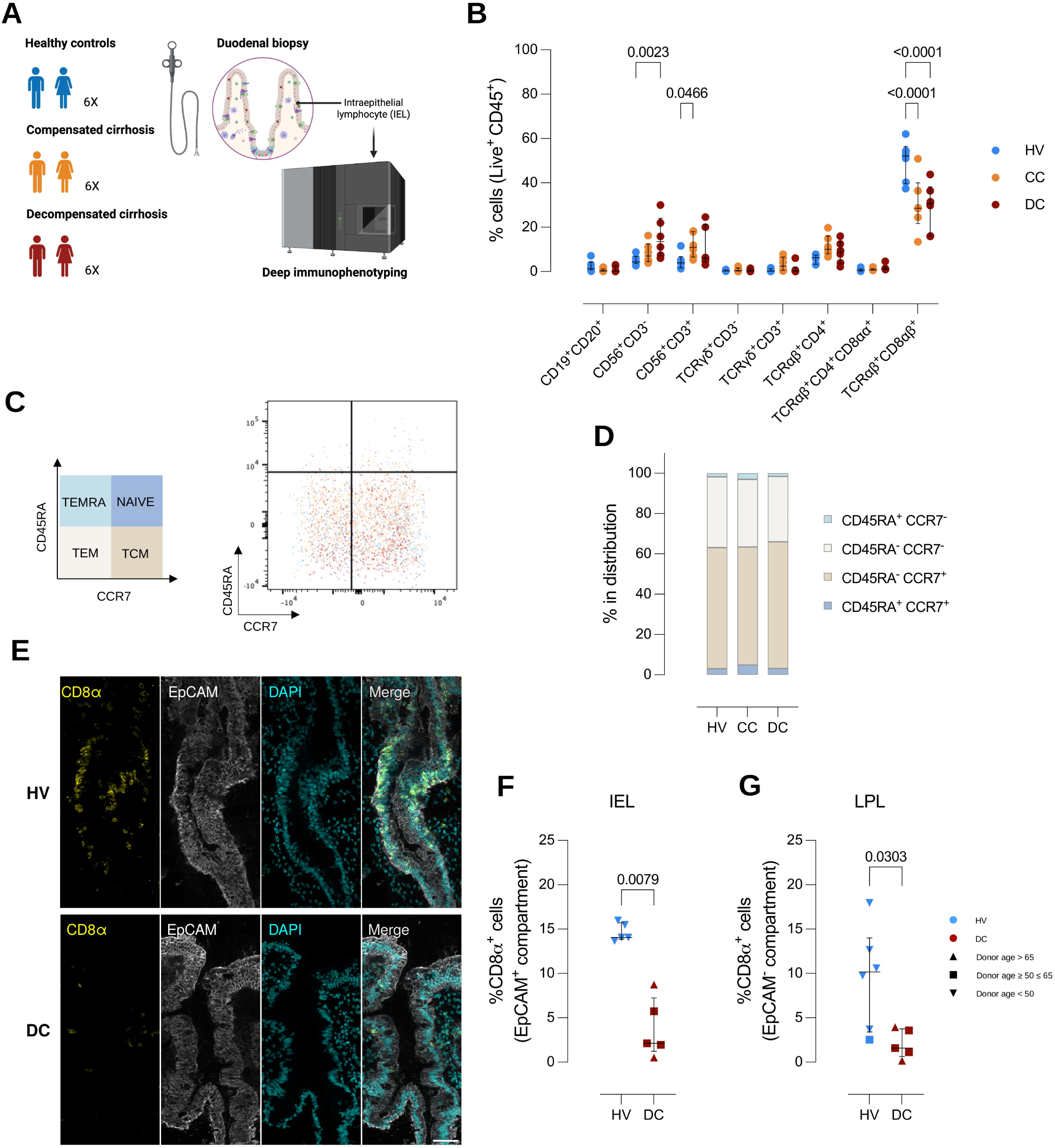
The intestinal epithelial immune compartment undergoes marked remodeling in cirrhosis. **(A)** Study design: deep immunophenotyping of duodenal IELs from HV, CC, and DC patients. **(B)** Frequency of major IEL subsets among live CD45⁺ cells, including CD19⁺CD20⁺ B cells, CD56⁺CD3⁻ NK cells, CD56^+^CD3^+^ NKT cells, and T-cell populations (TCRγδ⁺CD3⁻, TCRγδ⁺CD3⁺, TCRαβ⁺CD4⁺, TCRαβ⁺CD4⁺CD8αα⁺, and TCRαβ⁺CD8αβ⁺) across HV (n = 6), CC (n = 6) and DC (n = 6). **(C)** Definition (left) and representative flow cytometry plots (right) of CD8⁺ T cells in IEL differentiation states based on CD45RA and CCR7 expression: naïve (CD45RA⁺CCR7⁺), central memory (TCM, CD45RA⁻CCR7⁺), effector memory (TEM, CD45RA⁻CCR7⁻), and TEMRA (CD45RA⁺CCR7⁻), overlaid for HV (blue), CC (orange), and DC (red). **(D)** Distribution of CD8⁺ T cells in IEL subsets (naïve, TCM, TEM, TEMRA) shown as stacked bar plots. **(E)** Representative confocal photomicrographies of CD8α, EpCAM, and DAPI staining on duodenum tissue slices from HV and DC. Scale bar = 50 µm. **(F,G)** Quantification of CD8α^+^ cell frequencies within the **(F)** intraepithelial (EpCAM^+^) and **(G)** lamina propria (EpCAM^-^) compartments on duodenum tissue slices from HV (n = 6) and DC (n = 5). Data are presented as median ± IQR. **(F,G)** Outlier, if present, was removed with Grubbs’ method (alpha = 0.05). Statistical analysis was performed with **(B)** a two-way ANOVA and Bonferroni’s multiple comparison test comparing CC and DC with HV or **(F,G)** a Mann-Whitney test; p values ≤0.05 are indicated, symbols are coded by donor age.

NKT cells and NK cells were significantly expanded in compensated and decompensated cirrhosis respectively, whereas CD8αβ⁺ T cells, the dominant adaptive immune population in the healthy epithelium, were markedly reduced in both compensated and decompensated disease. Other IEL subsets remained comparatively stable. Importantly, analysis of CCR7/CD45RA-defined differentiation states demonstrated that most IELs maintained central and effector memory phenotypes irrespective of disease status (Fig. 5C,D), indicating that loss of CD8αβ⁺ IELs was not explained by major shifts in differentiation state.

To validate epithelial CD8⁺ T-cell depletion in situ, duodenal tissue sections were stained for CD8α and EpCAM (Fig. 5E). Quantitative image analysis confirmed significant reduction of CD8α⁺ T cells within the epithelial compartment of decompensated cirrhosis compared with healthy controls (Fig. 5F). A similar reduction was observed within the lamina propria compartment (Fig. 5G), indicating coordinated depletion of intestinal CD8⁺ T cells across mucosal compartments.

Collectively, these findings identify selective loss of epithelial CD8αβ⁺ T-cell surveillance as a major feature of mucosal immune remodeling in cirrhosis.

### Loss of intestinal CD8αβ⁺ T cells is associated with coordinated CCR9 dysregulation and markers of bacterial translocation

Because CCR9-dependent trafficking is central to small intestinal localization of CD8αβ⁺ T cells (Stenstad *et al*., 2007), we next investigated whether altered gut-homing programs could account for defective mucosal immune compartmentalization in cirrhosis.

The frequency of α4β7⁺ and αEβ7⁺ CD8αβ⁺ T cells, integrins associated with intestinal trafficking and epithelial retention, respectively, remained comparable across groups (Fig. 6A). In contrast, CCR9⁺CD8αβ⁺ T cells were profoundly reduced in cirrhosis, with approximately fivefold lower frequencies compared with healthy volunteers (Fig. 6B,C).

**Fig. 6.**
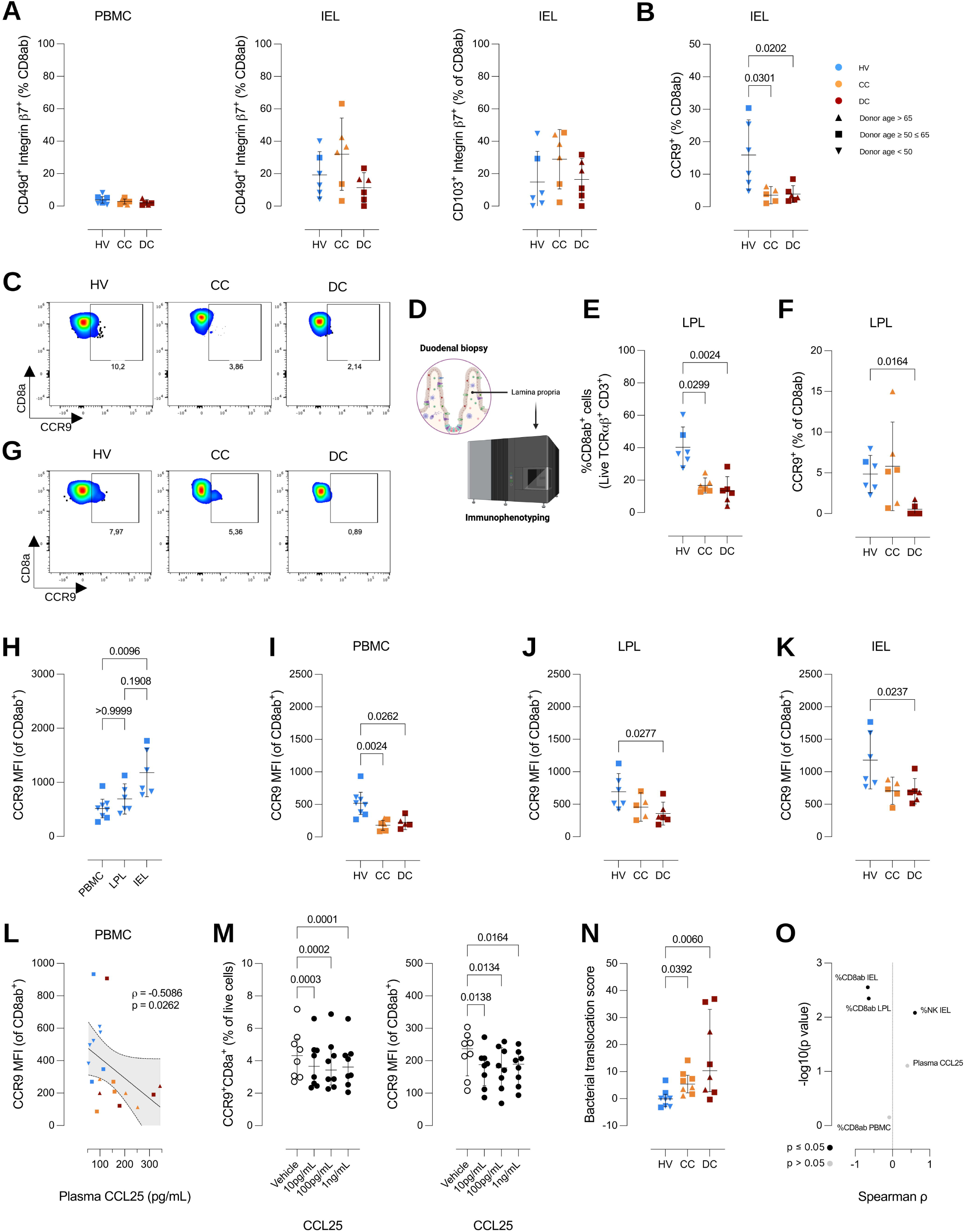
Loss of intestinal CD8αβ⁺ T cells is associated with coordinated CCR9 dysregulation. **(A)** Expression of gut-and epithelial-homing markers α4β7 on CD8αβ⁺ PBMCs and IELs, and αEβ7 on CD8αβ⁺ IELs, shown as percentage of parent cells. **(B)** Expression of small intestinal epithelium homing marker CCR9 on CD8αβ⁺ IELs, shown as percentage of parent cells. **(C)** Representative plots of CD8αβ⁺ IELs across groups; numbers indicate frequencies of parent populations. **(D)** Study design: deep immunophenotyping of duodenal lamina propria lymphocytes (LPLs) from the same cohorts. **(E)** Frequency of CD8αβ⁺ T LPLs among live TCRαβ⁺CD3⁺ cells. **(F)** Proportion of CCR9⁺ cells within CD8αβ⁺ T LPLs. **(G)** Representative plots for CD8⁺ T LPLs across groups; numbers indicate frequencies of parent populations. **(H)** CCR9 mean fluorescence intensity (MFI) in TCRαβ⁺CD8αβ⁺ T cells from HV across PBMC (n = 8), LPL (n = 6) and IEL (n = 6) compartments. **(I)** CCR9 MFI in circulating TCRαβ⁺CD8αβ⁺ T cells across HV (n = 8), CC (n = 7), and DC (n = 6). **(J)** CCR9 MFI in TCRαβ⁺CD8αβ⁺ T LPLs across groups. **(K)** CCR9 MFI in TCRαβ⁺CD8αβ⁺ T IELs across groups. **(L)** Consolidated Spearman correlation of CCL25 plasma concentrations and CCR9 MFI across HV (n = 8), CC (n = 6), and DC (n = 5). **(M)** CCR9 frequency and MFI after treatment of PBMCs from HVs (n = 8) with recombinant human CCL25 in concentrations spanning healthy and patient plasma CCL25 concentrations. **(N)** Consolidated EndoCab IgA, IgG and IgM, and lipopolysaccharide binding protein (LBP) ELISA assays into a single score reflecting bacterial translocation from HV (n = 8), CC (n = 8) and DC (n = 8). **(O)** Spearman correlations of the bacterial translocation score and the annotated patient parameters. Each dot represents one donor. Data are presented as median ± IQR. Outlier, if present, was removed with Grubbs’ method (alpha = 0.05). (**A, B, E, F, J, K)** HV (n = 6), CC (n = 6), and DC (n = 6). **(A,B,E,F,H-K,N)** Statistical analysis was performed with a Kruskal-Wallis test and Dunn’s multiple comparisons test comparing CC or DC with HV, **(M)** or with a repeated-measures one-way ANOVA with assumption of sphericity and Tukey’s multiple comparisons test; p values ≤0.05 are indicated, symbols are coded by donor age.

Because intestinal lymphocytes transit through the lamina propria before entering the epithelial layer, we next evaluated the lamina propria compartment (Fig. 6D). Consistent with intraepithelial findings, lamina propria CD8αβ⁺ T-cell frequencies were significantly reduced in cirrhosis and were similarly associated with loss of CCR9⁺CD8αβ⁺ T cells (Fig. 6E–G).

Assessment of CCR9 surface expression further revealed a physiological compartmental gradient in healthy individuals, with lowest expression in blood and highest expression within intraepithelial CD8αβ⁺ T cells (Fig. 6H). Across all compartments, cirrhosis was associated with a coordinated reduction of CCR9 surface expression, with the strongest decrease observed in decompensated patients (Fig. 6I–K). These findings were not driven by age and remained unchanged in sensitivity analyses in which hepatocellular carcinoma-positive donors were excluded (Fig. EV4).

Sustained ligand exposure drives chemokine receptor desensitization and internalization (Lamme *et al*, 2025). Chronically elevated systemic CCL25 would thus be expected to reduce functional surface CCR9. Consistently, there was a significant negative correlation between plasma CCL25 concentration and CCR9 surface expression on peripheral blood mononuclear cells (PBMCs) (Fig. 6L). Moreover, exposure of PBMCs from HVs to recombinant CCL25 at concentrations spanning those measured in healthy and patient plasma significantly reduced cell-surface CCR9 (Fig. 6M). Chronic systemic CCL25 elevation is therefore mechanistically consistent with, rather than contradictory to, a reduction in intestinal CD8αβ⁺ T-cells.

Furthermore, to assess the consequences of the loss of intraepithelial CD8αβ⁺ T-cells, we analyzed markers associated with bacterial translocation. Protein concentrations of EndoCab IgA, IgG and IgM, and lipopolysaccharide binding protein (LBP) were determined in plasma and consolidated into a single score reflecting bacterial translocation, which was significantly higher in disease compared with HV (Fig. 6N, Fig. EV5A-D). Additionally, there was a significant negative correlation between the bacterial translocation score and %CD8αβ⁺ T-cells in the intraepithelial and lamina propria compartments (Fig. 6O, Fig. EV5E,F), which exemplifies the concurrent loss of CD8αβ⁺ T-cells and an increase in systemic markers associated with bacterial translocation.

Together, these findings identify coordinated disruption of the CCL25–CCR9 axis across blood, lamina propria, and epithelial compartments and support defective intestinal immune compartmentalization as a central feature of cirrhosis-associated immune dysfunction.

### Single-cell RNA sequencing reveals antimicrobial attenuation and innate-like cytotoxic remodeling of intraepithelial lymphocytes

To define transcriptional alterations underlying intraepithelial immune remodeling, we initially performed single-cell RNA sequencing (scRNA-seq) of duodenal IELs isolated from healthy volunteers and patients with compensated cirrhosis. After quality control, 3,968 lymphocytes and 384 epithelial cells were retained for analysis (Appendix Fig. S3A,B). This initial dataset provided an overview of the transcriptional landscape of duodenal IELs and revealed features consistent with immune remodeling in cirrhosis (Appendix Fig. S3C). We subsequently expanded the scRNA-seq cohort to enable a more comprehensive characterization across disease stages and robust donor-level analyses. The expanded cohort comprised 19 donors, including healthy volunteers and patients with compensated and decompensated cirrhosis, with healthy donors ranging in age from 22–68 years. 132,386 cells were retained after quality control, including 28,460 IELs and 103,926 epithelial cells, for downstream analyses.

Unsupervised clustering identified six principal immune populations, including two CD8αβ⁺ T-cell states, a CD4⁺ T-cell subset, a TCRγδ^+^CD3^+^ T-cell subset, a mature and a cycling innate subsets (Fig. 7A,B). Consistent with flow cytometry findings, CD8αβ⁺ T-cell populations dominated the epithelial immune compartment.

**Fig. 7.**
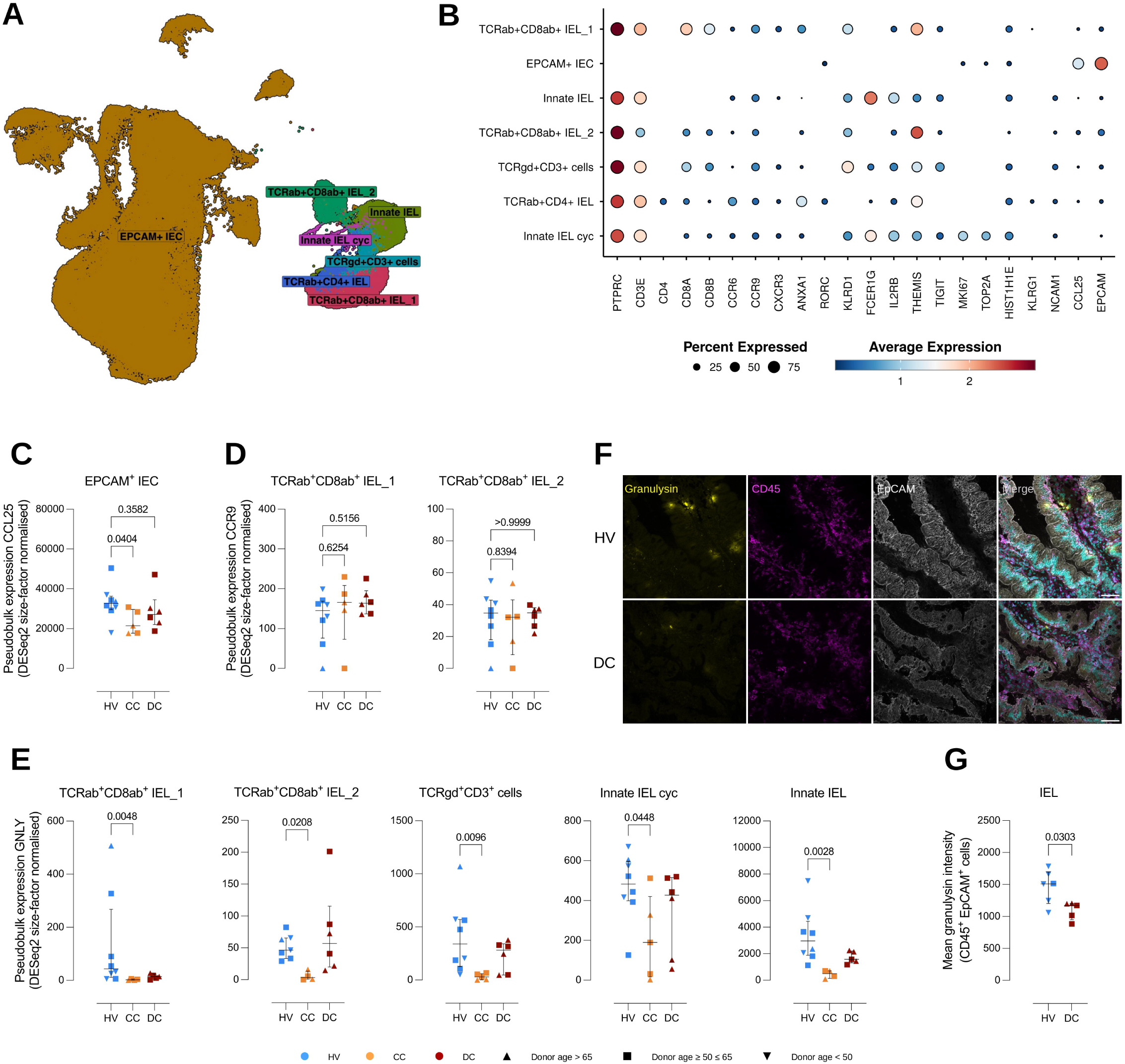
Single-cell RNA sequencing reveals antimicrobial attenuation and cytotoxic remodeling of intraepithelial lymphocytes. **(A)** UMAP embedding of single-cell transcriptomes showing major IEL populations alongside EPCAM⁺ epithelial cells. **(B)** Dot plot summarizing canonical marker genes defining IEL subsets and intestinal epithelial cells (IEC). **(C)** Pseudobulk expression of CCL25 in the EPCAM^+^ cell cluster across groups. **(D)** Pseudobulk expression of CCR9 in TCRαβ^+^CD8αβ^+^_1 and TCRαβ^+^CD8αβ^+^_2 cell clusters across groups. **(E)** Pseudobulk expression of GNLY in different immune populations across groups. **(F)** Representative photomicrographies of granulysin, CD45, and EpCAM staining. Scale bar = 50µm. **(G)** Quantification of granulysin staining in HV (n = 6) and DC (n = 5). Data are presented as median ± IQR. Outlier, if present, was removed with Grubbs’ method (alpha = 0.05). Statistical analysis was performed with **(C,D,E)** a Kruskal-Wallis test and Dunn’s multiple comparisons test, or **(G)** a Mann-Whitney test. p values ≤0.05 are indicated, symbols are coded by donor age.

We next interrogated the CCL25–CCR9 axis at the transcriptional level. CCL25 expression was restricted to epithelial cells, whereas CCR9 expression was predominantly detected within lymphocyte populations (Fig. EV5G). Despite persistent systemic elevation of circulating CCL25 protein, epithelial *CCL25* transcription was not increased in cirrhosis (Fig. 7C). Likewise, *CCR9* transcript abundance remained largely preserved across CD8αβ⁺ T-cell populations (Fig. 7D). These findings indicate that reduced CCR9 surface expression in cirrhosis is unlikely to result from transcriptional repression and instead support post-transcriptional dysregulation of the receptor.

Differential expression analysis identified two principal features of IEL remodeling in cirrhosis. First, antimicrobial defense programs were attenuated across lymphocyte populations, including a reduced expression of *GNLY* (Fig. 7E). This was conserved on protein level (Fig. 7F,G). Additionally, in an exploratory analysis, granulysin protein expression correlated with the bacterial translocation score (Fig. EV5H). Second, residual IELs exhibited enhanced cytotoxic transcriptional programs characterized by increased granzyme expression, including *GZMA* and *GZMH* (Fig. EV5I,J). All group-level statistics were computed on per-donor pseudobulk aggregates.

To determine whether these transcriptional changes reflected convergent clonal expansion, we performed paired TCR repertoire analysis. IEL repertoires were highly oligoclonal in both healthy volunteers and cirrhosis patients, with dominant clonotypes occupying a large fraction of the compartment (Appendix Fig. S3D). Expanded clonotypes were largely private to individual donors, and no consistent disease-associated TCR convergence was observed (Appendix Fig. S3E,F).

Collectively, single-cell analysis identifies qualitative remodeling of the residual IEL compartment in cirrhosis characterized by attenuation of antimicrobial programs together with enrichment of cytotoxic pathways, while supporting non-transcriptional dysregulation of the CCL25–CCR9 axis.

## Discussion

Bacterial infections are major drivers of decompensation and mortality in cirrhosis, placing immune dysfunction at the center of disease progression. Cirrhosis-associated immune dysfunction is traditionally conceptualized as the coexistence of systemic inflammation and immune paralysis, frequently extending to adaptive compartments through the framework of T-cell exhaustion (Noor & Manoria, 2017; Yadav *et al*, 2025). Here, integrated profiling of paired blood and duodenal mucosa instead identifies a compartmentalized form of immune dysfunction, characterized by preserved systemic lymphocyte competence but defective intestinal immune deployment.

We found no evidence of bona fide T-cell exhaustion. Classical exhaustion is defined by sustained inhibitory receptor expression together with progressive functional impairment (Barber *et al*., 2006; Blank *et al*., 2019; Wherry, 2011), yet neither circulating nor mucosal T cells showed enrichment of PD-1 or CTLA-4, and circulating lymphocytes retained robust cytokine responsiveness despite progressive inflammatory remodeling. This is consistent with recent studies suggesting that exhaustion phenotypes emerge predominantly during acute-on-chronic liver failure rather than in baseline cirrhosis (Liang *et al*, 2026; Yao *et al*, 2024; Yu *et al*, 2024). Circulating B-cell frequencies and subsets were likewise preserved, in apparent contrast with the B-cell depletion reported in the liver in hepatitis B virus-related cirrhosis (Bai *et al*, 2023). Because these studies differ in tissue compartment, etiology and methodology, the two observations are compatible, and together support the view that immune remodeling in cirrhosis is compartment-specific rather than uniform.

In contrast to the preserved systemic compartment, the intestinal epithelial immune interface was markedly reshaped. The healthy duodenal epithelium was dominated by CD8αβ⁺ IELs, whereas cirrhosis was associated with their depletion and with expansion of innate cytotoxic populations. Conventional CD8αβ⁺ IELs play central roles in epithelial surveillance and microbial containment (Huleatt *et al*, 2001; Masopust *et al*, 2001a; Masopust *et al*, 2001b; Sheridan *et al*, 2014), and the expansion of innate cytotoxic populations resembles remodeling associated with epithelial injury and barrier dysfunction in inflammatory mucosal diseases (Calleja *et al*, 2011; Schmitz *et al*, 2016; Uhde *et al*, 2020). The residual IELs were also qualitatively altered, with attenuated antimicrobial programs and enriched cytotoxic pathways at transcriptional level. Reduced granulysin, confirmed at the protein level, points to impaired intracellular antimicrobial defense (Walch *et al*, 2014), whereas increased granzyme expression indicates enhanced cytotoxic differentiation (Chawla *et al*, 2025; Meresse *et al*, 2006). This imbalance may compromise pathogen containment while promoting epithelial injury.

Consistent with impaired barrier defense, a composite score of plasma markers associated with bacterial translocation (endotoxin core antibodies and lipopolysaccharide-binding protein) was elevated in cirrhosis and correlated inversely with CD8αβ⁺ T-cell frequencies in both the epithelium and lamina propria. In an exploratory analysis, it also correlated inversely with tissue granulysin protein. These associative correlations are consistent with a compromised barrier integrity accompanying IEL loss at a major site of microbial translocation in cirrhosis.

Our data identify dysregulation of the CCL25–CCR9 axis as a candidate mechanism for this defective compartmentalization. CCR9 is a key regulator of small intestinal T-cell homing (Stenstad *et al*., 2007), and cirrhosis was associated with a coordinated reduction of surface CCR9 on CD8αβ⁺ T cells in blood, lamina propria and epithelium. This occurred despite preserved α4β7 and αEβ7 integrin expression, indicating selective disruption of CCR9-dependent trafficking rather than a global loss of gut-homing capacity. Preserved *CCR9* transcript abundance in IELs further points to post-transcriptional regulation of the receptor.

The coexistence of elevated plasma CCL25 with reduced intraepithelial CD8αβ⁺ T cells may appear paradoxical. However, IEL recruitment is governed by the local epithelial CCL25 gradient rather than by the systemic pool (Stenstad *et al*., 2007), and epithelial *CCL25* transcription was not increased in cirrhosis. Systemic and mucosal CCL25 are therefore dissociated. Because sustained chemokine exposure drives receptor desensitization and internalization (Lamme *et al*., 2025), chronically elevated circulating CCL25, already detectable in compensated cirrhosis, would be expected to reduce functional surface CCR9 before cells reach the intestine. Indeed, exposure of healthy-donor PBMCs to CCL25 at concentrations spanning those measured in patient plasma reduced surface CCR9, providing direct experimental support for this mechanism. Given that aberrant CCL25 expression has been reported in chronic liver disease (Eksteen *et al*, 2004; Graham *et al*, 2022), the diseased liver may represent a source of this systemic chemokine. CCL25 was prioritized for mechanistic follow-up because it is the only mediator altered in cirrhosis with a defined homing receptor on the cells under investigation; other mediators, such as hepatocyte growth factor, may nonetheless carry prognostic information (Table EV1).

Several limitations warrant consideration. The cross-sectional design precludes temporal or causal inference and the limited yield of human intraepithelial lymphocytes precluded migration assays, so impaired intestinal homing remains to be validated. Cohort and limited paired tissue-plasma samples restricted etiological stratification and correlation analyses. Nevertheless, the concordant findings across orthogonal platforms and tissue compartments support the robustness of the observed immune remodeling.

Healthy volunteers were younger than patients, an imbalance intrinsic to cirrhosis, which rarely presents before middle age. However, age did not correlate significantly with any readout within the cirrhosis groups, and compensated patients were older yet displayed the milder phenotype. Overall, effect sizes were preserved when patients were compared only with the three healthy volunteers within the patient age range, and the oldest healthy volunteer contributing duodenal tissue displayed the most preserved CCR9 phenotype. Excluding donors with hepatocellular carcinoma did not alter statistical significance. Nevertheless, a fully age-matched duodenal biopsy cohort would provide the most direct control.

Collectively, our findings support a revised framework in which cirrhosis-associated immune dysfunction reflects defective spatial organization of adaptive immunity rather than generalized immune suppression: systemic lymphocytes remain competent but fail to establish effective epithelial surveillance at the intestinal barrier. Restoring mucosal immune compartmentalization, for instance by targeting the CCL25–CCR9 axis, may therefore represent a therapeutic avenue distinct from, and complementary to, global immune enhancement.

## Material and Methods

### Patients and study design

Healthy volunteers and patients with compensated and decompensated liver cirrhosis were prospectively recruited at the University Hospital UZ Leuven. All participants provided written informed consent prior to inclusion, and experiments conformed to the Declaration of Helsinki and the Department of Health and Human Services Belmont Report. The study was approved by the Medical Ethics Committee of the University Hospital Leuven (reference numbers: S50354, S65781 and S68855).

Healthy controls had no history or clinical evidence of chronic liver disease and no features of portal hypertension. Cirrhosis was defined according to established clinical, biochemical, and/or radiological criteria. Disease severity was assessed using the Model for End-Stage Liver Disease (MELD) score and Child–Pugh classification (A5–C15). Patients were systematically characterized for complications of portal hypertension, including ascites, gastroesophageal varices, hepatorenal syndrome, and hepatocellular carcinoma, as well as for disease etiology, including metabolic dysfunction–associated steatotic liver disease (MetALD/MASH) and portal vein thrombosis. Compensated and decompensated cirrhosis were distinguished according to the Baveno VII framework (de Franchis *et al*, 2022), where transition from the compensated to the decompensated state is defined by the development of ascites, variceal hemorrhage, or overt hepatic encephalopathy. Compensated cirrhosis was defined by the absence of all these complications.

Demographic and anthropometric data (age, sex, body mass index) and relevant comorbidities (diabetes mellitus, arterial hypertension, dyslipidemia, and chronic kidney disease) were recorded. No restrictions were applied regarding age or sex.

Participants were excluded if they had a history of liver transplantation or colectomy, were pregnant or breastfeeding, or had active HIV infection. To limit confounding of mucosal immune and microbiome-related readouts, individuals receiving systemic immunosuppressive therapy, or with prior fecal microbiota transplantation were excluded. Subjects with known or suspected chronic gastrointestinal disorders, including inflammatory bowel disease, irritable bowel syndrome, or celiac disease, were also excluded due to their established impact on epithelial and immune homeostasis. No participant in any group had received antibiotics in the period preceding sampling. Participation in an interventional clinical trial, or the presence of any severe or uncontrolled comorbidity or concomitant treatment deemed by the investigators to compromise safety or data integrity, also led to exclusion.

### Blood and duodenal mucosa biopsy processing

#### Isolation of intraepithelial lymphocytes (IEL)

This protocol was optimized to enrich for IEL and was used to obtain cells for spectral flow cytometry and 5′ scRNA-seq. Briefly, six biopsies per individual were immediately collected into a 15-mL tube containing 10 mL of ice-cold chelating solution composed of 5.6 mM Na₂HPO₄, 8.0 mM KH₂PO₄, 96.2 mM RNase-free NaCl, 1.6 mM RNase-free KCl, 43.4 mM D-sucrose, 54.9 mM D-sorbitol, and 0.5 mM dithiothreitol (DTT), prepared in DNase/RNase-free ultrapure water. Biopsies were washed three times with ten successive rinses in this solution. The solution was then replaced with 10 mL of chelating solution supplemented with 10 mM ethylenediaminetetraacetic acid (EDTA). Tubes were placed on a HulaMixer platform at 4°C and gently agitated for 45 min. Following incubation, six consecutive 10-mL fractions containing epithelial crypts were collected, pooled, and divided into two 50-mL conical tubes containing 10 mL of dialyzed fetal bovine serum (FBS) to neutralize residual EDTA. The pooled crypt suspension was centrifuged at 100 x g for 2 min at 4°C, resulting in a pellet enriched for epithelial crypts and a supernatant containing IEL. The supernatant containing IEL was collected, filtered through a 70-µm cell strainer, and centrifuged at 500 x g for 5 min at 4°C. The pellet was resuspended in 100 µL of FACS Wash (2.5% FBS, 2 mM EDTA in 1X PBS), and IEL were counted using Trypan Blue exclusion.

#### Generation of a single-cell suspension of duodenal epithelial cells and IEL

This protocol was designed to recover intestinal epithelial cells, together with IEL that remain associated with the crypt fraction, for 3′ scRNA-seq. Duodenal biopsies were processed as described above up to the 100 × g centrifugation step. The crypt-enriched pellet was then resuspended in 3 mL of TrypLE Select (10x diluted to 5x in D-PBS) and 10 µg/mL DNAse I (pre-warmed to 37 °C). Enzymatic digestion was performed for up to 3 minutes at 37 °C, aided by mechanical dissociation using a 1 mL pipette, with one pipetting cycle every 2 seconds. To terminate digestion, 7 mL of D-PBS (without Ca²⁺/Mg²⁺) supplemented with 10 mM EDTA, 2% dialyzed FBS, and 12.5 mM HEPES was added. The resulting cell suspension was passed through a 40 µm Flowmi filter to remove debris and undigested material, followed by three wash steps consisting of centrifugation at 500 × g for 5 minutes and resuspension in 10 mL of fresh buffer. Viable cells were counted, and the suspension was adjusted to a final concentration of 1,000 cells/µL. Cells were immediately processed for scRNA-seq as described below.

#### Isolation of lamina propria lymphocytes (LPL)

LPLs were isolated as previously described (Kim *et al*, 2022). Briefly, tissue biopsies were digested for 35 minutes at 37 °C. The resulting cell suspension was then washed, filtered, and subjected to density gradient centrifugation using a 40/80% Percoll solution. Lymphocytes collected from the interface were thoroughly washed before proceeding to staining procedures.

#### Isolation of peripheral blood mononuclear cells (PBMC) and plasma

Blood was collected in EDTA-coated tubes and processed within 1 hour of collection. PBMCs were purified by density gradient centrifugation using Lymphoprep according to the manufacturer’s instructions. Isolated cells were cryopreserved in FBS containing 10% dimethyl sulfoxide (DMSO) and stored in liquid nitrogen until further use. Plasma was extracted by centrifugation (300 x *g*, 20 minutes, 20°C) and stored at -80°C until use.

### Spectral flow cytometry staining

Patient PBMCs (2 x10^6^ cells/well), LPLs, and IELs were seeded in V-bottom 96-well plates as paired unstained and fully stained wells per compartment. Single-stained controls consisted of buffy coat PBMCs (2 x 10^6^ cells/well) spiked with patient cells along with UltraComp eBeads Compensation Beads (1 drop/well). Unless otherwise stated, centrifugation was at 600 x g for 5 min at 4°C, with cells washed in FACS Buffer #1 (1% FBS, 2 mM EDTA in 1X PBS) and beads in 1X PBS. Cells were blocked with 2.4G2 Fc block (1:1000 in PBS, 30 min, 4°C), washed, and stained with viability dye (Zombie NIR) in 1X PBS (20 min at room temperature in the dark). Unstained and single-cell controls were processed in parallel without viability dye. Surface staining was performed in Brilliant Buffer with 20% 2.4G2 Fc block) for 1 h at room temperature in the dark and washed once with FACS Buffer #1. Cells were fixed with Foxp3/Transcription Factor Staining Buffer Set (30 min, 4°C), washed twice with 1X Permeabilization Buffer, and stained intracellularly in 1X Permeabilization Buffer with 20% 2.4G2 Fc block (overnight, 4°C). Samples were washed twice with 1X Permeabilization Buffer and once with FACS Buffer #1, transferred to 1.2 mL polypropylene deep-well plates and stored at 4°C in the dark until acquisition on a Sony ID7000 flow cytometer equipped with 355, 405, 488, 561, and 640 nm lasers.

Antibodies against the following markers were used: CD49a/Integrin α1, CD3, CD2, IgG, CD56/NCAM, IgM, Ki67, CD8β, FoxP3, CD4, CD127/IL7Rα, CD1c, CD14, CD15, CD33, CD34, CD123, CD203c, CD27, HLA-ABC, CD45RA, T-bet, EpCAM/CD326, CD103/Integrin αE, GATA3, CD45, CD8α, CD161, TCRαβ, FcεRI, CD19, IgD, IgA, CD69, CD20, RORγt, CTLA4/CD152, CD49d/Integrin α4, TCRγδ, CD45RO, KLRG1, CD160, CCR7/CD197, Integrin β7, CD38, PD-1/CD279, CCR9/CD199, and HLA-DR.

### Immunofluorescence staining and image analysis

Small intestinal pinch biopsies were embedded in OCT for cryosectioning. 10 µm thick sections were used for immunofluorescence staining. In short, sections were fixed with 4% PFA at room temperature for 30 min, after which they were incubated in block buffer (5% normal donkey serum, 3% bovine serum albumin, 0.1% Triton X-100 in 1x TBS) for 2 h at room temperature (RT). Primary antibody incubation was performed in block buffer overnight at 4°C. Slides were washed with TBS and incubated with secondary antibody for 1.5 h at RT in the dark. Subsequently, slides were washed with TBS, stained with 5 µg/mL DAPI in TBS for 10 min, and mounted. Images were acquired on a Zeiss LSM 780 or Zeiss LSM 980 (25x and 10x objective respectively). Images were imported in ImageJ and analysed using the CLIJ2 package (Haase *et al*, 2020). DAPI was used to segment individual cells using Voronoi-Otsu labeling. Mean intensities of the separate channels were recorded in individual cells, after which an absolute threshold was used to differentiate CD8α^+^ T cells or CD45^+^ lymphocytes within the EpCAM^-^ (lamina propria) and EpCAM^+^ (intraepithelial) compartments. The fraction of CD8α^+^ T cells was determined by dividing the number of positive cells by the total number of cells in the respective compartment.

### T cell activation assay

Flat-bottom 96-well plates were coated overnight at 4°C with anti-CD3 antibody (eBioscience, clone OKT3) at 2 µg/mL in 1X PBS. Control wells received PBS only. Equal cell numbers were seeded into anti-CD3–coated wells supplemented with anti-CD28 antibody (5 µg/mL; activated condition) or into PBS-coated wells (non-activated condition). After 20 h of incubation at 37°C in 5% CO₂, 50 µL of conditioned medium were collected per well for Olink Target 48 Cytokine and Immune Surveillance profiling. GolgiPlug (Brefeldin A, 1:1000) was then added, and cells were incubated for an additional 4 h at 37°C.

### Flow cytometry staining for intracellular cytokines

Following supernatant collection, cells were transferred to V-bottom 96-well plates and centrifuged (400 × g, 5 min, 4°C); supernatants were stored at −20°C. Cells were washed in FACS Buffer #2 (2.5% FBS, 2 mM EDTA in 1× PBS), and surface staining was performed as described above until fixation step. Fixation followed the “Dish Soap Protocol” (Burton *et al*, 2025). Briefly, cells and beads were fixed with fixative (2% formaldehyde, 0.05% Dreft (or Fairy), 0.5% Tween-20, 0.1% Triton X-100 in 1X PBS) for 30 min at room temperature in the dark. Samples were centrifuged at 600 *g* for 5 min at 4°C and permeabilized in Perm Buffer (0.05% Dreft in 1X PBS) for 30 min at room temperature in the dark, washed twice with FACS Buffer #2 (beads with 1X PBS), and stained overnight intracellularly in FACS Buffer #2 containing 20% 2.4G2 Fc block at 4°C in the dark. Cells were then washed twice with FACS Buffer #2, transferred to 1.2-mL deep-well plates, and stored at 4°C in the dark until acquisition on the Sony ID7000 spectral cell analyzer.

Antibodies against the following markers were used: CD4, CD3, CD56, TNFα, IFNγ, IL-10, IL-1β, CD14, IL-4, IL-17A, T-bet, TCR Vα7.2, GATA3, CD8α, CD19, IL-6, RORγt, TCRγδ, CXCL8, IL-22, FoxP3, Granzyme A, Granzyme B, TCRαβ, CD28, and viability dye.

Pair-wise comparisons were performed using R software via paired t-test comparing the percentage of intracellular-protein positive cells within each cellular subset, unstimulated vs. stimulated, within each patient group (HV, CC, DC). For this purpose, the rstatix package was used. Data filtering and manipulation were performed using the Tidyverse package, and heatmaps were created using the Morpheus tool.

### Spectral flow cytometry unmixing and data analysis

Data were acquired on a Sony ID7000 spectral cell analyzer, and fluorescence was collected as full emission spectra across all detectors for each laser line. Single-stained controls (PBMC spiked with tissue cells or beads) for each fluorochrome and an unstained control were used to generate reference spectra, which were then applied for spectral unmixing using the instrument’s weighted least-squares algorithm implemented in the ID7000 software (Sony Biotechnology). For highly auto-fluorescent samples, distinct autofluorescence signatures were identified from unstained cells using the Autofluorescence Finder tool and incorporated as additional reference spectra during unmixing to improve separation of dim populations. The resulting unmixed parameters were exported as compensated fluorescence intensities and used for all downstream gating and analysis with FlowJo 10.10.

### Olink proteomics analysis

Protein levels were quantified on plasma samples and on PBMC conditioned media using the Olink® Target 48 Cytokine and Immune Surveillance panels following the manufacturer’s instructions. Olink data were processed using the Tidyverse package in R. Proteins whose expression did not appear in at least 5 samples were removed, and the remaining null values were estimated using the k-nearest neighbors’ algorithm, present in the R impute package. Differentially expressed proteins were determined using the Kruskal-Wallis test, with Dunn’s post-hoc test, available in the FSA package. Additionally, the heatmap was created using the Morpheus tool.

### Functional assays

#### CCR9 desensitization

Buffy coat-derived PBMCs were thawed and cultured in basal medium (GlutaMAX supplemented RPMI-1640, 1x HEPES, 1x Antibiotic-Antimycotic, 1x sodium pyruvate, and 1x MEM non-essential amino acids) supplemented with 10% FBS, for 24 h at 37°C, 5% CO_2_ before starting any functional assays. Additionally, PBMCs were equilibrated in assay medium (basal medium supplemented with 0.5% BSA) for 1 h before starting any functional assays.

PBMCs were exposed to a range of physiologically relevant recombinant CCL25 concentrations for 1 h in assay medium, after which live cell surface staining for spectral flow cytometry was performed using antibodies against the following markers: CD8α, CCR9/CD199, CD3, CD4, CD19, CD56/NCAM, CD14. Data were acquired on an Aurora spectral flow cytometer (Cytek Biosciences). Unmixing was performed with the SpectroFlo software using the weighted least squares algorithm. Manual gating and quality control were performed using FlowJo.

### ELISA assays

ELISA assays were performed according to manufacturer’s instructions. Plasma samples were diluted 1000x (LBP), 400x (IgG), or 50x (IgA, IgM). Additionally, a BCA protein assay was performed to allow for normalization of samples, for which plasma was diluted 100x. Standards and samples were run in triplicate, and measured on a Molecular Devices SpectraMax iD3 (BCA), or on a BioTek 800 TS (LBP, IgA, IgG, IgM) at the wavelength recommended by the manufacturer. OD values from the standards were plotted, and a 4-parameter logistic curve was fit. Sample concentration was determined by interpolation on this curve, normalized based on their dilution, and further normalized based on total protein content as determined by the BCA assay. ELISA assays were used to calculate a robust Z-score reflecting bacterial translocation by summation of factors that increase, and subtraction of factors that decrease in disease compared with HV.

### Single-cell RNA sequencing and bioinformatics analysis

#### Single-cell RNA sequencing

Following preparation of IEL single-cell suspensions for spectral flow cytometry, whenever sufficient cells were available, an aliquot of viable cells was immediately processed at the Laboratory of Mucosal Biology for single-cell RNA sequencing using the 10x Genomics Chromium platform (Chromium GEM-X Single Cell 5′ Chip v3, PN #1000698; Chromium GEM-X Single Cell 5′ Kit v3, PN #1000699). Gene expression (GEX) libraries were generated using the Chromium GEM-X Single Cell 5′ Library Construction Kit C (PN #1000694), and paired T cell receptor (TCR) libraries were generated using the Human TCR Amplification Kit (PN #1000252), according to the manufacturer’s instructions. Additionally, following digestion of crypts to obtain single-cell suspensions containing epithelial cells and IEL, viable cells were immediately processed for single-cell RNA sequencing using the 10x Genomics Chromium platform (Chromium GEM-X 3’ chip v4, PN #1000690; Chromium GEM-X Single Cell 3’ Kit v4, PN #1000691; Chromium Next GEM Single Cell 3’ kit v3.1, PN #1000268; Chromium Next GEM Chip G Single Cell Kit PN #1000127).

Sequencing was performed at the KU Leuven Genomics Core using an Illumina NextSeq 2000 platform with SP 100-cycle flow cells. Paired-end sequencing was conducted with the following configuration: Read 1 (28 bp), i7 index (8 bp), and Read 2 (91 bp).

Reads were aligned to the human reference genome (GRCh38, GRCh38-3.0.0) using the Cell Ranger count pipeline, which incorporates the STAR aligner (Dobin *et al*, 2013). Gene-barcode matrices were subsequently imported into R (v4.3.2) for downstream analysis using Seurat (v5.5.0) (Hao *et al*, 2021).

#### Ambient RNA correction

Ambient RNA contamination was estimated and corrected using SoupX (Young & Behjati, 2020). Ambient RNA profiles were inferred from droplets with low UMI counts, and a contamination fraction was estimated for each cell. Gene expression counts were then adjusted based on the inferred ambient RNA profile, and corrected count matrices were used for all downstream analyses.

#### Preprocessing, integration, and clustering

Quality control was performed to exclude low-quality cells and potential doublets. Cells with fewer than 500 detected genes, more than 6,000 detected genes, or >20% mitochondrial gene content were removed. Additional computational doublet detection was not performed on the 5’ samples due to the limited number of recovered cells, and stringent filtering thresholds were applied to mitigate potential doublet inclusion. The doubletFinder algorithm (McGinnis *et al*, 2019) was applied per sample on the 3’ samples using the first 20 principal components, a pN of 0.25, and a pK of 0.01. For each sample, the expected number of doublets was estimated as a ceiling proportion of each sample’s cell count. Doublet classifications and pANN scores were then assigned back to the corresponding cells in the full merged object. For each sample, counts were log-normalized using Seurat’s NormalizeData function, and highly variable features were identified using FindVariableFeatures. Data were scaled (ScaleData) and dimensionality reduction was performed using principal component analysis (PCA) using 30 principal components.

To account for inter-sample variability and batch effects across donors and conditions, datasets were integrated using Harmony (Korsunsky *et al*, 2019) on the PCA embeddings. For downstream analyses, 35 principal components were retained based on variance explained and dataset complexity.

A shared nearest neighbor graph was constructed using FindNeighbors (k.param = 30, method = “rann”), followed by clustering using FindClusters across a range of resolutions (0.4–3). Clustering stability and biological interpretability were assessed across resolutions, and clusters were merged where appropriate. Uniform Manifold Approximation and Projection (UMAP) was computed using RunUMAP based on the selected principal components for visualization.

#### Cell annotation strategy

Cell type and state annotation was performed using a multi-step approach integrating unsupervised clustering, canonical marker gene expression, and reference-based validation. Cluster identities were initially inferred by examining the top differentially expressed genes (Wilcoxon rank-sum test) across clusters and resolutions.

Major immune lineages were assigned based on established marker genes. Annotations were cross-referenced against publicly available human intestinal single-cell atlases using multiple platforms, including CellxGene (Chan Zuckerberg Initiative), the Cell Annotation Platform, and the Broad Institute Single Cell Portal. This integrative strategy ensured consistency with established transcriptional identities while preserving dataset-specific resolution. Upon cell annotation, clusters corresponding to non-epithelial/non-lymphoid cells (mast cells and macrophages) were excluded prior to downstream analysis.

#### Differential gene expression

Genes were considered significant at adjusted p <0.05, with stringent thresholds (log₂FC > 1 and adjusted p <0.05) applied for downstream analyses and visualization. Pseudobulk count matrices were normalized using the DESeq2 algorithm before comparing gene expression between donors. Interpretation of transcriptional programs (e.g., antimicrobial or cytotoxic) was based on concordant changes across curated gene sets.

#### TCR clonotype analysis

TCR libraries were processed using the Cell Ranger V(D)J pipeline with default parameters and the GRCh38-vdj reference (v7.1.0). Productive TCR sequences were defined by in-frame CDR3 regions without stop codons. Cells with at least one productive TCR chain were retained. Clonotypes were defined based on identical CDR3 amino acid sequences of the TCR β chain, incorporating paired α chains where available, and analyzed using the immunarch package (ImmunoMind, 2019).

Clonal expansion was quantified as the frequency of each clonotype within a sample, and repertoire structure was assessed using clonotype frequency distributions and cumulative abundance of dominant clones. Oligoclonality was defined by the proportion of cells occupied by the most expanded clonotypes (e.g., top 1 and top 10). Analyses were performed on a per-sample basis to account for differences in cell yield. Clonotype overlap between samples was assessed using pairwise CDR3-aa clonotype-sharing analysis. Under our current experimental conditions, expanded clonotypes were largely sample-specific, with no cross-sample sharing.

### Statistical analysis

Statistical analyses were performed using GraphPad Prism v10.4.0 (GraphPad Software), unless stated otherwise. Parametric or non-parametric tests with post-hoc multiple comparisons (corrected for multiple testing) were applied as appropriate; specific tests are detailed in the figure legends. All p-values are two-sided, with significance set at p <0.05. Exact sample sizes, statistical tests and p-values are stated in each figure legend.

## Data availabilit*y*

Raw sequencing data have been deposited in the European Genome-Phenome Archive (EGA) under accession numbers EGAS50000001717, EGAS50000002194, and EGAS50000002169. Source data underlying the figures are provided with this paper. Raw imaging data are available on the BioImage Archive under accession number S-BIAD4162. All other data supporting the findings of this study are available from the corresponding author upon reasonable request. Source data for all main figures have been deposited in BioStudies and are linked to the corresponding figures in the online version of this article. [BioStudies accession to be added upon assignment].

## Code availability statement

Code used for Olink analysis is available at: https://github.com/alex-denadai/Olink.

## Disclosure and competing interest statement

The authors declare that they have no known competing financial interests or personal relationships that could have appeared to influence the work reported in this paper.

## Acknowledgments

We thank Drs. Hanne Hellinckx, Ingrid Demedts, Timothy Adam, Jan Vandewalle, Charlotte Van Laeken, Hanne Vandeputte, and Prof. Benedicte Dubois for their valuable assistance with clinical sample collection. We gratefully acknowledge Dr. Hannelie Korf (KU Leuven) for her critical revision of the manuscript. We also thank the Genomics Core Leuven (KU Leuven), and the Cell and Tissue Imaging Cluster; Zeiss LSM 780 – SP Mai Tai HP DS, supported by Hercules AKUL/11/37 and FWO G.0929.15 to Pieter Vanden Berghe, University of Leuven; and Zeiss LSM 980 –Insight X3 multiphoton microscope Supported by FWO, Multi-Phosthar, I001122N and FBI-LIAISE, I000123N, Cell and Tissue Imaging Cluster (CIC), Pieter Vanden Berghe, KU Leuven, University of Leuven. We would like to thank Prof. Koen Poesen and Amir Ghodrati for their support with ELISA assays.

## Funding

A.D.-S. was supported by a KU Leuven start-up grant (STG/22/023) and a KU Leuven C1 grant (C14/23/135). TV is supported by FWO through a senior clinical research mandate (1830517N).

## Figure Expanded View (EV) Captions

**Figure EV1.**
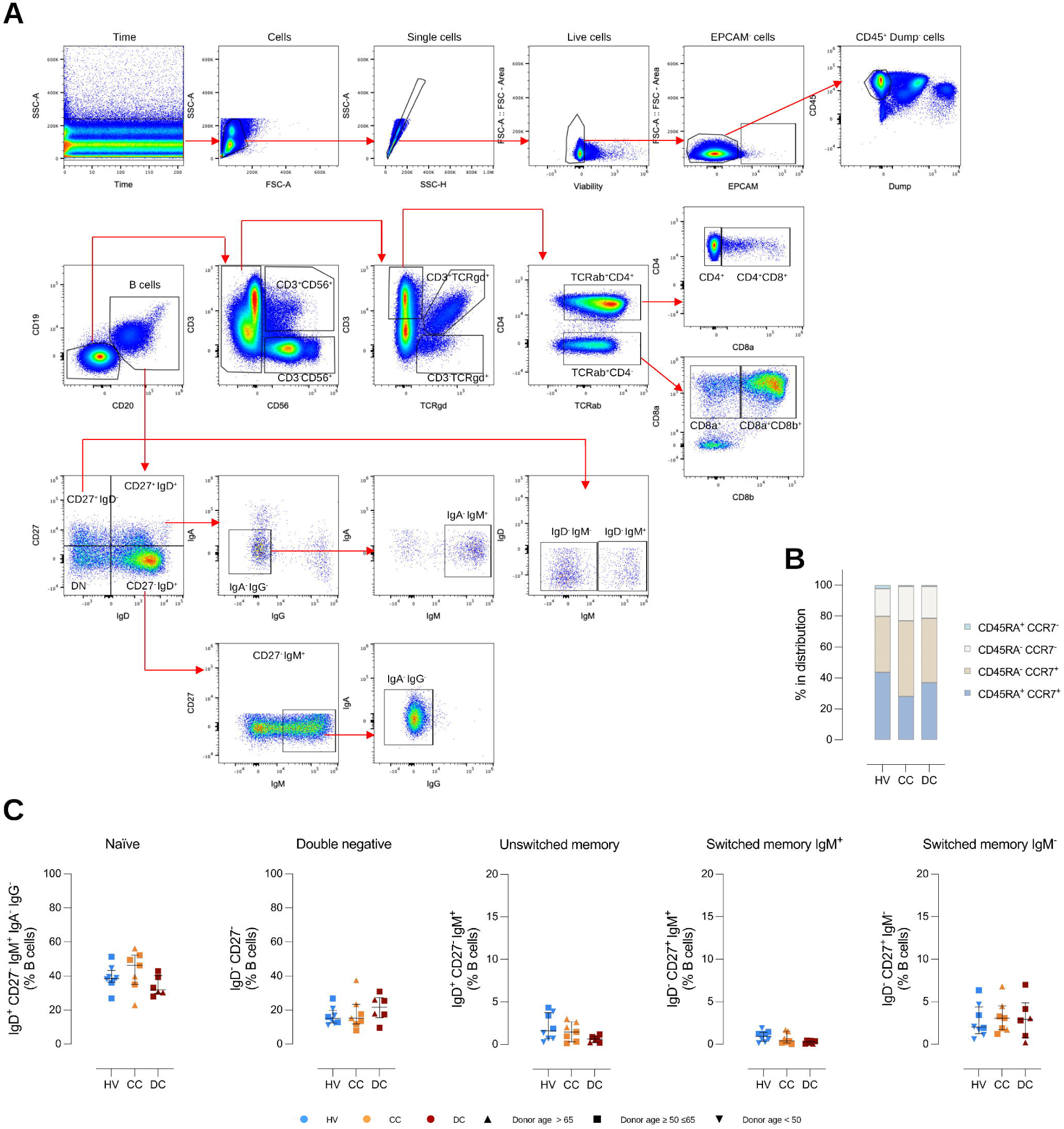
Gating strategy for high-dimensional immunophenotyping. **(A)** Representative flow cytometry gating strategy for identification of immune cell subsets in PBMCs. After exclusion of acquisition irregularities, cells were defined based on FSC-A versus SSC-A, followed by doublet exclusion using SSC-A versus SSC-H. Live cells were selected by exclusion of viability dye–positive events, and epithelial cells were excluded by gating out EPCAM⁺ cells. CD45⁺ leukocytes were identified after removal of lineage-positive dump-negative channel events. B cells were defined as CD19⁺CD20⁺ and further subdivided into naïve and class-switched populations based on IgD, IgM, IgG, and IgA expression. NK and NKT cells were identified as CD3⁻CD56⁺ and CD3⁺CD56⁺, respectively. CD3⁺ T cells were separated into TCRαβ⁺ and TCRγδ⁺ populations. Within TCRαβ⁺ cells, CD4⁺ and CD8⁺ T-cell subsets (CD8αα and CD8αβ) were defined. Red arrows indicate the sequential gating hierarchy. **(B)** Distribution of CD4⁺ T helper cells – naïve (CD45RA⁺CCR7⁺), central memory (CD45RA⁻CCR7⁺), effector memory (CD45RA⁻CCR7⁻), and TEMRA (CD45RA⁺CCR7⁻) – in PBMCs shown as stacked bar plots for each group. **(C)** B cell subsets – naïve, double negative, unswitched memory, switched memory IgM^+^, and switched memory IgM^-^ – across HV (n = 8), CC (n = 7), and DC (n = 6). **(C)** Data are presented as median ± IQR. Outlier, if present, was removed with Grubbs’ method (alpha = 0.05). Statistical analysis was performed with a Kruskal-Wallis test and Dunn’s multiple comparisons test, comparing CC and DC with HV; p values ≤0.05 are indicated, symbols are coded by donor age.

**Figure EV2.**
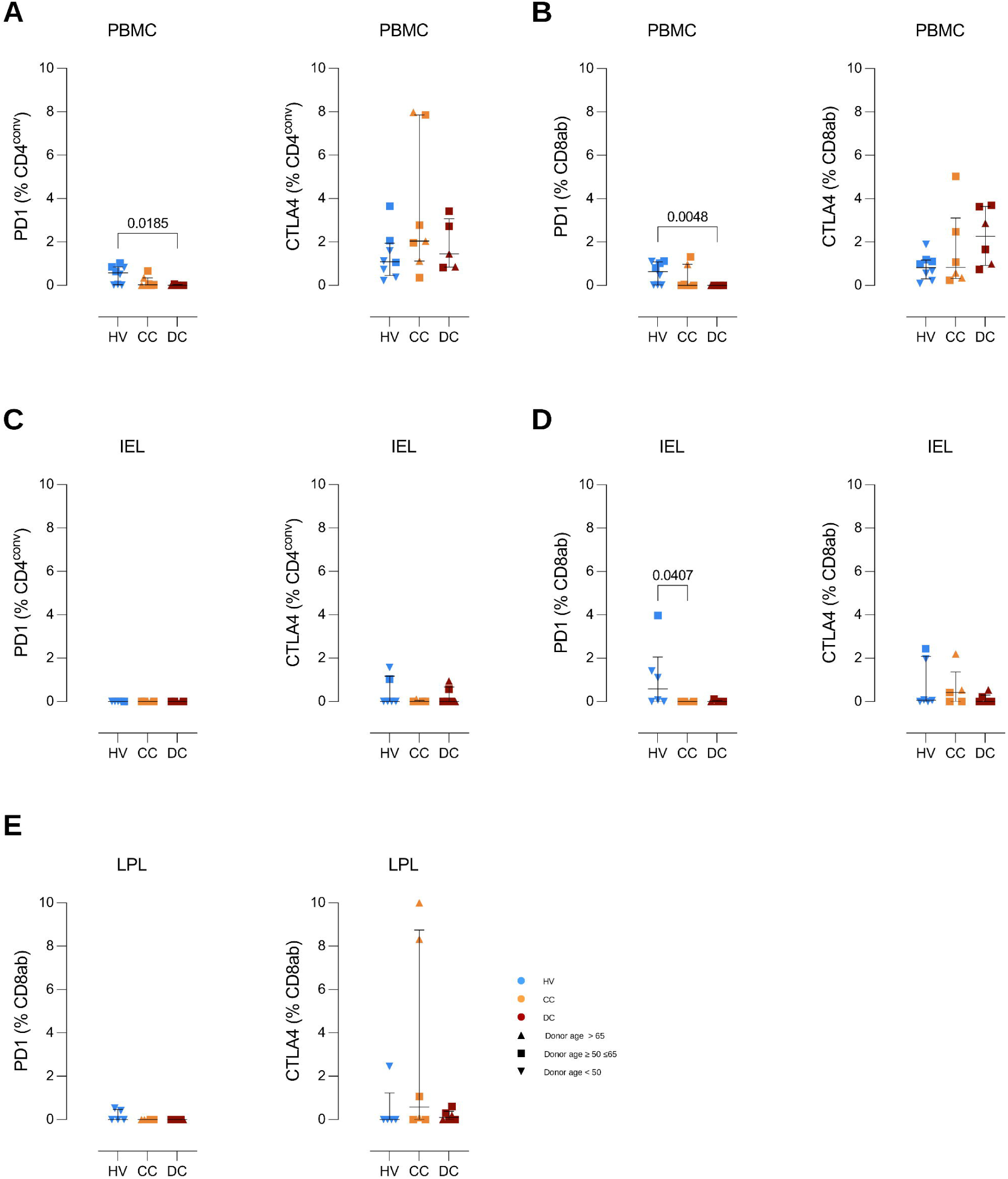
Canonical exhaustion markers are not upregulated in cirrhosis. **(A)** Frequency of PD-1^+^ and CTLA4^+^ cells in circulating conventional CD4⁺ T cells. **(B)** Frequency of PD-1^+^ and CTLA4^+^ cells in circulating conventional CD8αβ⁺ T cells. **(C)** Frequency of PD-1^+^ and CTLA4^+^ cells in CD4⁺ IEL. **(D)** Frequency of PD-1^+^ and CTLA4^+^ cells in CD8αβ⁺ IEL. **(E)** Frequency of PD-1^+^ and CTLA4^+^ cells in intestinal lamina propria CD8αβ⁺ T cells. Data are presented as median ± IQR. Outlier, if present, was removed with Grubbs’ method (alpha = 0.05). Statistical analysis was performed with a Kruskal-Wallis test and Dunn’s multiple comparisons test, comparing CC and DC with HV; p values ≤0.05 are indicated, symbols are coded by donor age.

**Figure EV3.**
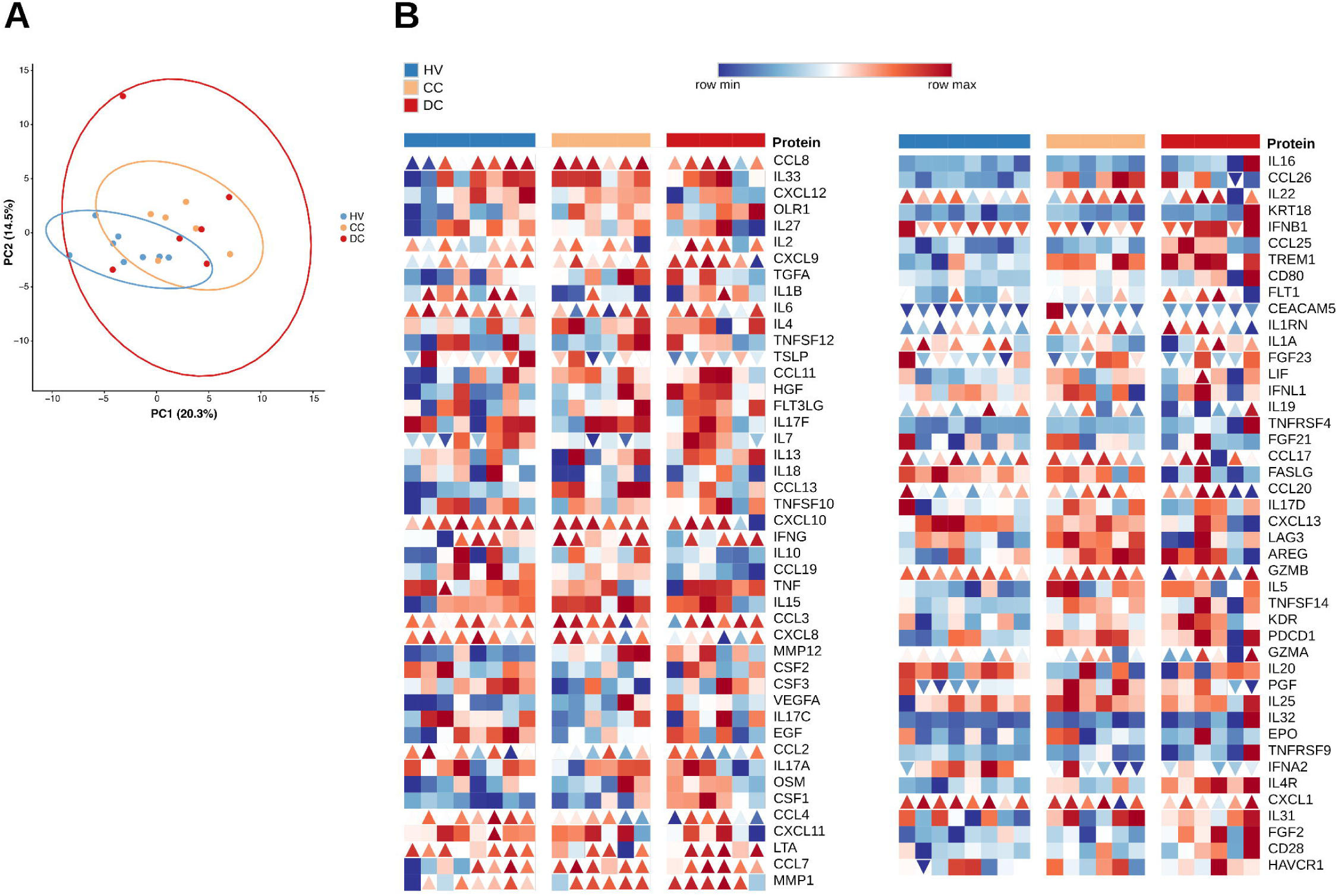
Preserved T cell effector function in cirrhosis. **(A)** Principal component analysis (PCA) of inflammation and immune proteins in supernatants from stimulated PBMCs. **(B)** Heatmap of inflammation and immune protein quantification in supernatants harvested from activated PBMCs. Scaled (z-score) protein levels are displayed. Triangles pointing upwards or downwards indicate proteins over or under assay detection limit, respectively. **(A,B)** HV (n = 8), CC (n = 6), and DC (n = 6). Statistical analysis was performed with a Kruskal-Wallis test and Dunn’s multiple comparisons test.

**Figure EV4.**
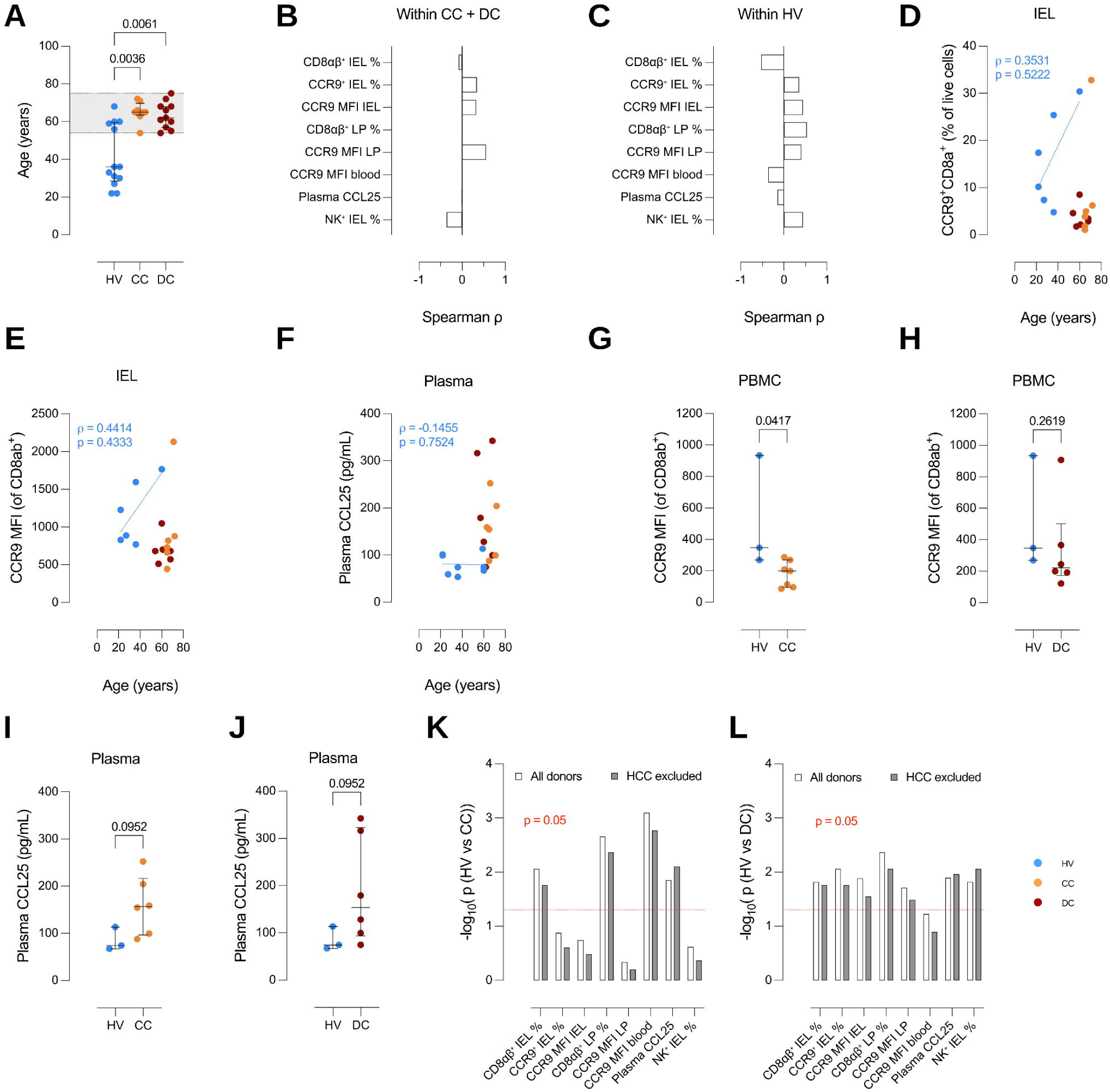
Age and hepatocellular carcinoma sensitivity analyses. **(A)** Age distribution of HV (n = 13), CC (n = 8) and DC (n = 11). Shading denotes the age range spanned by the patient groups. **(B,C)** Spearman correlation coefficients between age and each readout, computed **(B)** within the cirrhosis groups (CC (n = 7) and DC (n = 7) combined), **(C)** or in the HV group (n = 6) only; all p ≥ 0.067, all |ρ| ≤ 0.55. **(D–F)** Individual donor values for **(D)** CD8αβ⁺ IEL frequency, **(E)** CCR9 mean fluorescence intensity (MFI) in IEL, and **(F)** plasma CCL25 concentration plotted against donor age. Dashed lines show the least-squares fit within healthy volunteers; Spearman’s ρ and p-value are indicated for the groups. **(G,H)** Comparison of age-comparable healthy volunteers (≥ 50 years) for CCR9 MFI in blood with **(G)** compensated and **(H)** decompensated patients. **(I,J)** Comparison of age-comparable healthy volunteers (≥ 50 years) for plasma CCL25 with **(I)** compensated and **(J)** decompensated patients. **(K, L)** Effect of excluding the hepatocellular carcinoma (HCC)- positive donor in each patient group on **(K)** HV (n = 6) versus CC (n = 7) and **(L)** HV (n = 6) versus DC (n = 7) on the indicated readouts, shown as −log_10_(*p*) before and after exclusion; the dashed line marks p = 0.05. **(G – H)** HV (n = 3), CC (n = 7), DC (n = 6). **(I – J)** HV (n = 3), CC (n = 6), DC (n = 6). **(A,D-J)** Each dot represents one donor. **(A,G-J)** Statistical analysis was performed with a **(A)** Kruskal-Wallis test and Dunn’s multiple comparisons test, or a **(G-J)** Mann-Whitney test; all p values are indicated.

**Figure EV5.**
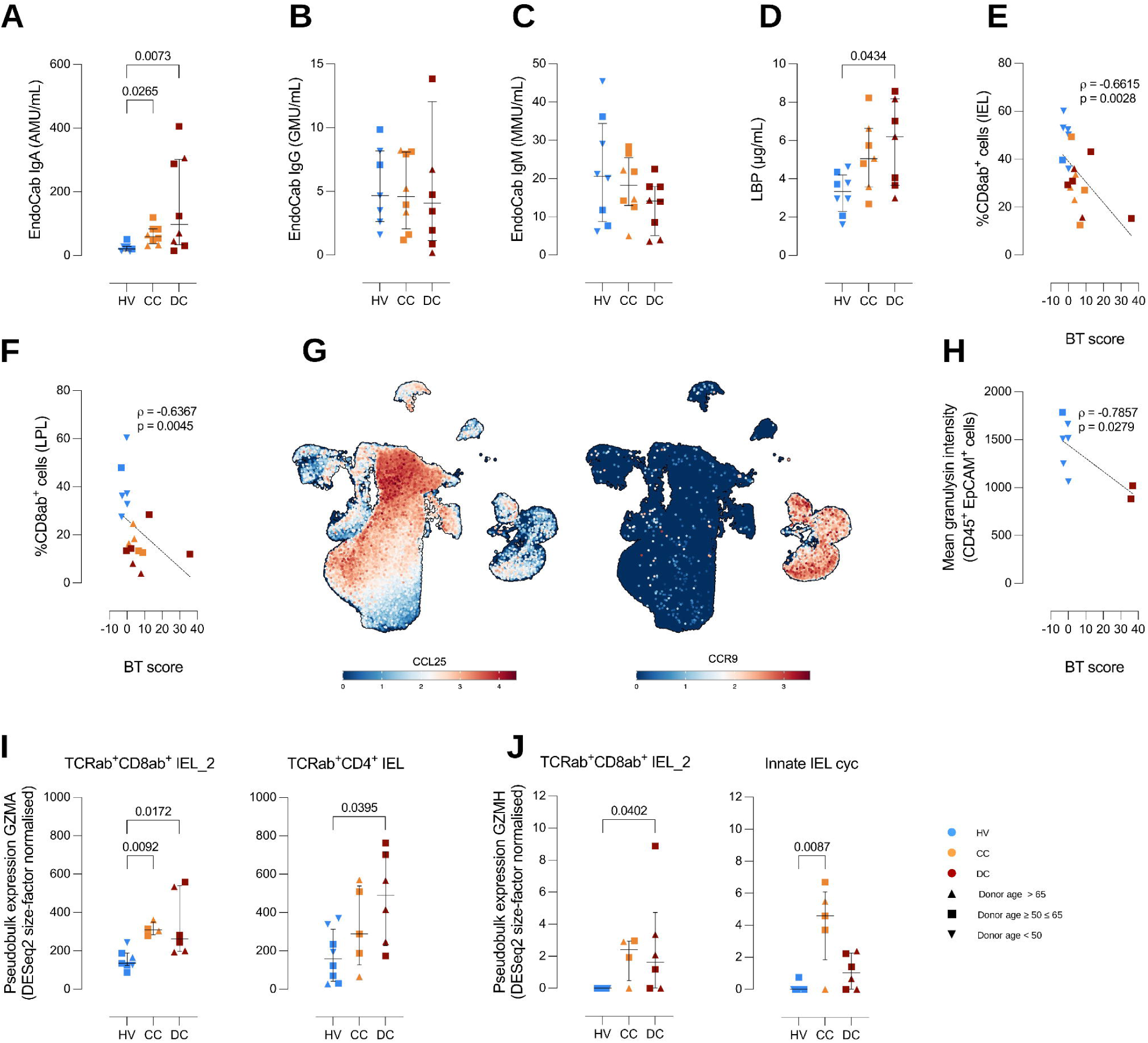
Bacterial translocation markers and qualitative changes in granzyme expression. (A-D) Plasma protein concentrations of (A) EndoCab IgA, (B) EndoCab IgG, **(C)** EndoCab IgM, and **(D)** lipopolysaccharide binding protein (LBP) in HV (n = 8), CC (n = 8), and DC (n = 8). **(E,F)** Spearman correlation of bacterial translocation (BT) score and %CD8ab^+^ cells in the **(E)** intraepithelial and **(F)** lamina propria compartments. **(G)** Feature plots showing the spatial distribution and relative expression of CCL25 and CCR9. **(H)** Spearman correlation of BT score and mean granulysin intensity from segmented IELs. **(I,J)** Pseudobulk expression of **(I)** *GZMA* and **(J)** *GZMH* in indicated cell clusters in HV (n = 8), CC (n = 5), and DC (n = 6). **(A-F,H-J)** Each dot represents one donor. **(A-D,I,J)** Data are presented as median ± IQR. Outlier, if present, was removed with Grubbs’ method (alpha = 0.05). Statistical analysis was performed with a Kruskal-Wallis test and Dunn’s multiple comparisons test, comparing CC and DC with HV; p values ≤0.05 are indicated, symbols are coded by donor age.

**Appendix Figure S1.**
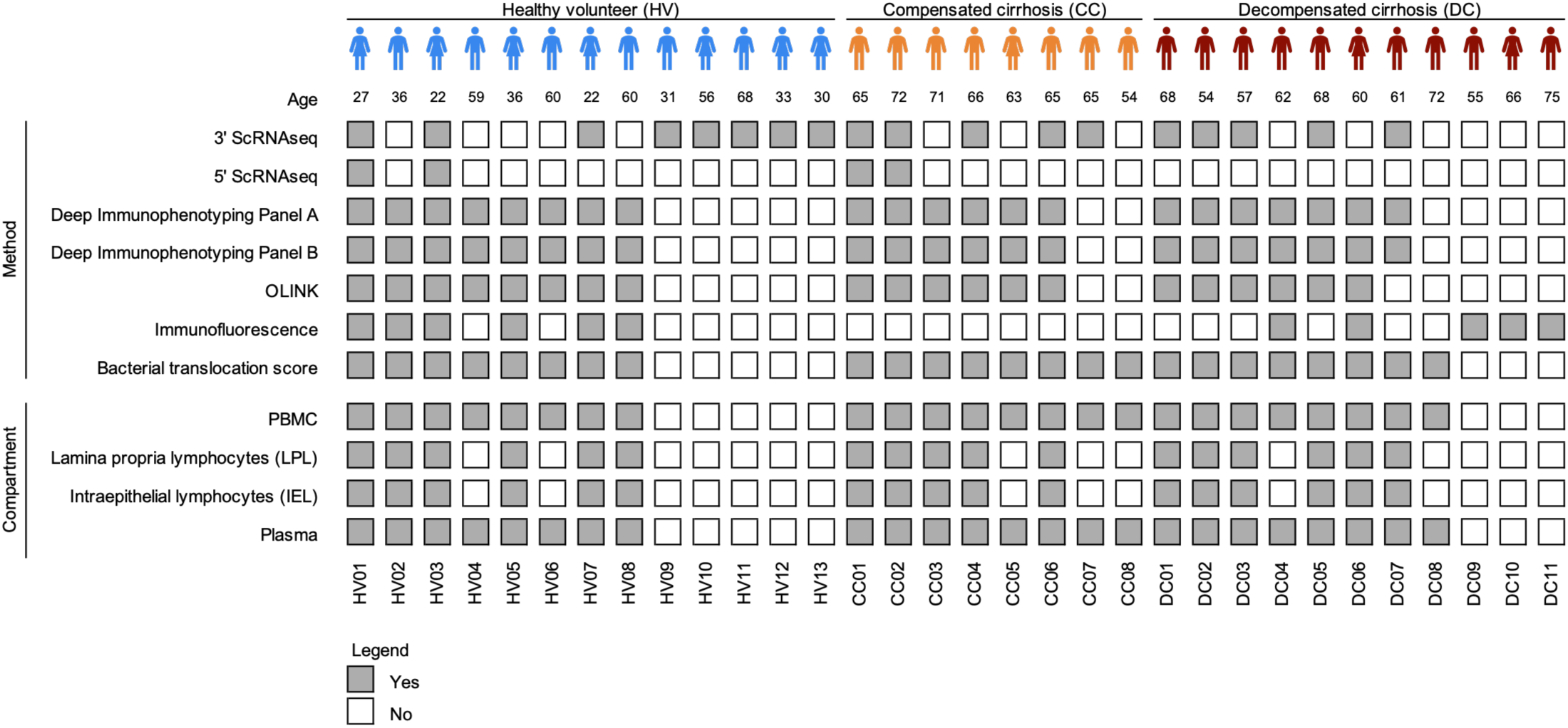
Overview of the human cohort and method allocation. Primary human cohort indicating the experimental approaches performed on each subject and the types of samples available per patient, including PBMCs, plasma, LPLs, and IELs.

**Appendix Figure S2.**
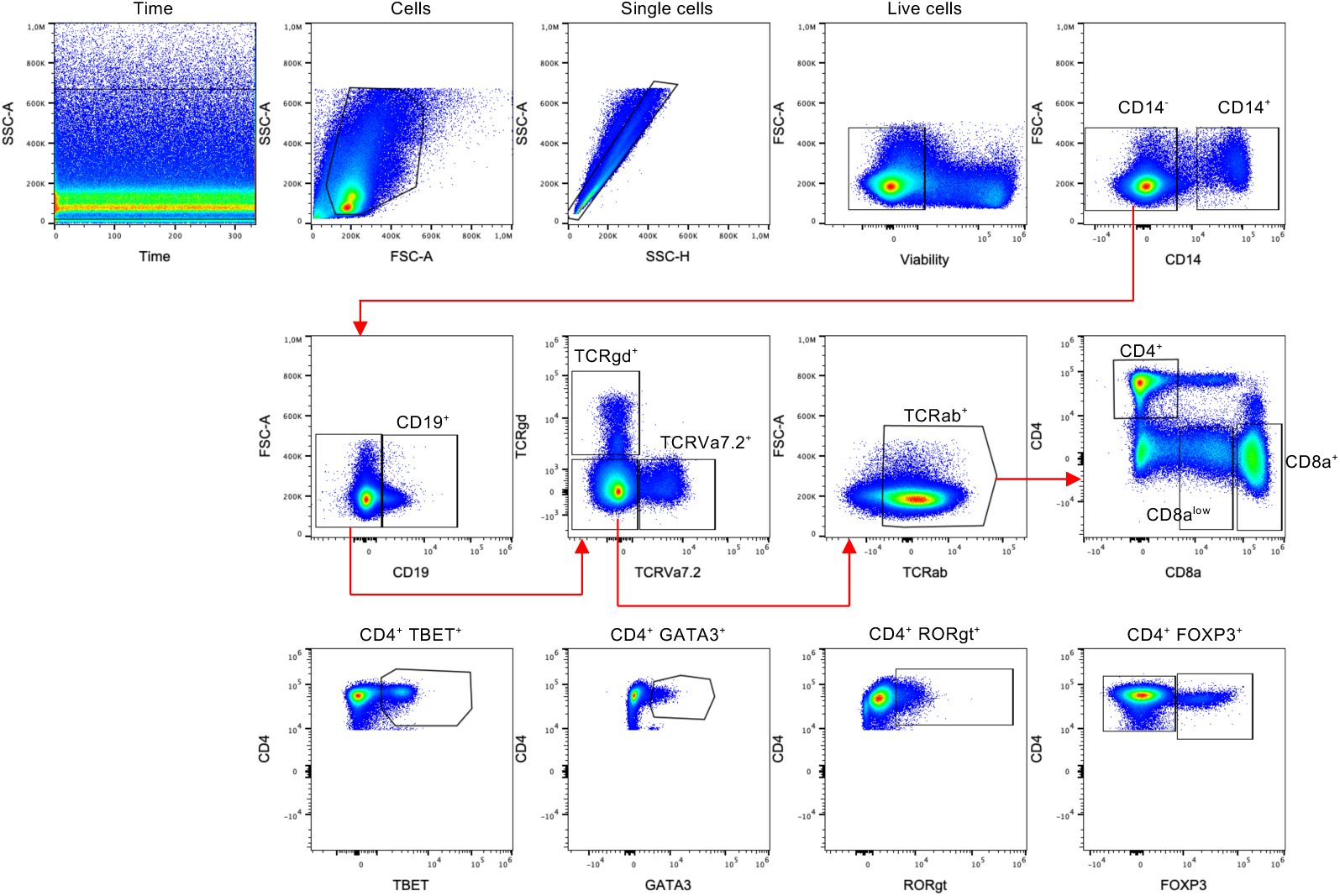
Gating strategy for assessment of intracellular cytokines. Representative flow cytometry gating strategy for identification of immune cell subsets in PBMCs. After exclusion of acquisition irregularities, cells were defined based on FSC-A versus SSC-A. Doublets were excluded using SSC-A versus SSC-H, and live cells were selected by exclusion of viability dye–positive events. Monocytes were identified as CD14⁺ cells, while CD14⁻ cells were carried forward for lymphocyte analysis. B cells were defined as CD19⁺. T cells were subdivided into TCRγδ⁺, TCRVα7.2⁺, and TCRαβ⁺ populations. TCRαβ⁺ cells were further separated into CD4⁺ and CD8α⁺ T cells. CD4⁺ T cells were characterized based on transcription factor expression, including TBET, GATA3, RORγt, and FOXP3, defining Th1, Th2, Th17, and regulatory T cell subsets, respectively. Red arrows indicate the sequential gating hierarchy.

**Appendix Figure S3.**
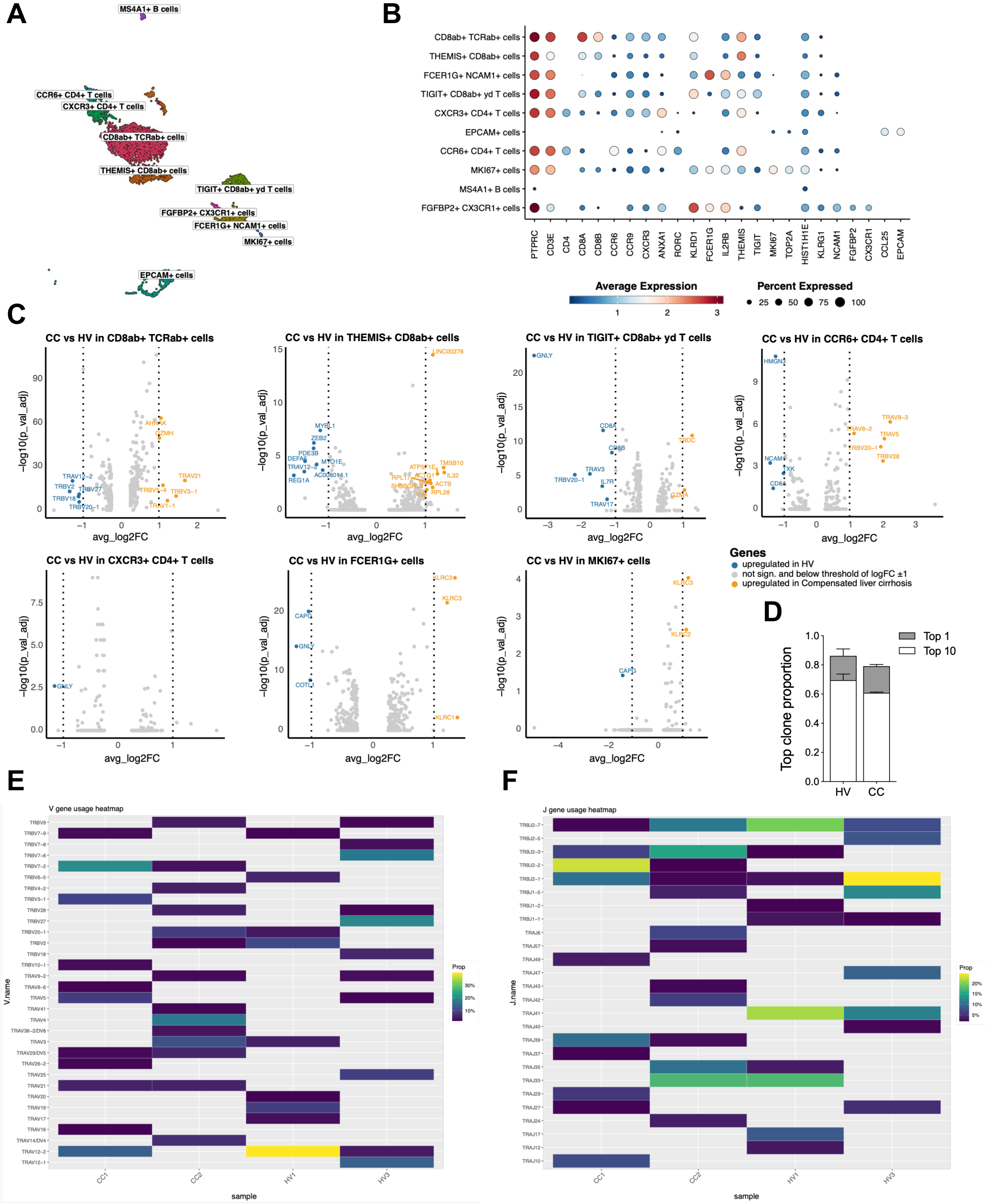
5’ ScRNA sequencing and V and J gene segment usage in TCRαβ repertoires across individual samples. **(A)** UMAP embedding of single-cell transcriptomes showing major IEL populations alongside EPCAM⁺ epithelial cells. **(B)** Dot plot summarizing canonical marker genes defining IEL subsets and intestinal epithelial cells (IEC). **(C)** Exploratory analysis illustrating differential gene expression within IEL subsets, highlighting significantly altered genes (log₂ fold change > 1, adjusted p < 0.05) between HV (n = 2) and CC (n = 2). **(D)** Bar plot depicting proportions of top 1 and top 10 TCRαβ clonotypes per group. Data is shown as median ± range. **(E)** Heatmap showing the relative frequency (%) of TCR V gene segment usage across individual samples from each study group. Each column represents a sample, and each row a distinct V gene segment. **(F)** Heatmap showing the relative frequency (%) of TCR J gene segment usage across the same samples, with identical layout.

## Notes

### Competing Interest Statement

The authors have declared no competing interest.

### Summary of Updates

This version has been substantially updated to strengthen the robustness of our findings and to extend them mechanistically and clinically. First, we addressed potential confounders in the patient cohort. Because healthy volunteers were on average younger than patients with cirrhosis, donor age is now indicated for every individual in all figures, and we examined the main differences within an age overlapping subset and in relation to age within each group. Key comparisons were also repeated after excluding patients with hepatocellular carcinoma to assess whether the observed changes are driven by malignancy. Second, we linked the loss of intestinal CD8 alpha beta intraepithelial lymphocytes to gut barrier failure. We measured circulating markers of bacterial translocation, namely endotoxin core antibodies and lipopolysaccharide binding protein, and combined them into a bacterial translocation score. This score correlates inversely with CD8 alpha beta T cell frequencies in the epithelium and lamina propria and with epithelial granulysin, connecting the depletion of these cells to the translocation of bacterial products that drives complications in cirrhosis. Third, we explored how these cells are lost. We extended the analysis of the CCL25 and CCR9 gut homing axis across blood, epithelium and lamina propria, measured plasma CCL25, and tested the effect of CCL25 on CCR9 expression in vitro. Together with the mapping of CCL25 to the epithelium and CCR9 to intraepithelial lymphocytes, these data implicate dysregulated gut homing in the depletion of intraepithelial lymphocytes in cirrhosis. Fourth, we added single cell RNA sequencing of duodenal biopsies. Pseudobulk analysis of intraepithelial lymphocyte subsets, together with immunofluorescence for granulysin, indicates that the remaining cells also show altered cytotoxic programmes, suggesting that both the number and the function of these protective cells are compromised. Finally, the immunophenotyping was expanded to B cell and memory T cell subsets and to the checkpoint molecules PD1 and CTLA4, the plasma proteomic analysis was presented in greater depth, and the text, methods, statistics and figure legends were revised throughout.

